# *MINPP1* prevents intracellular accumulation of the cation chelator inositol hexakisphosphate and is mutated in Pontocerebellar Hypoplasia

**DOI:** 10.1101/2020.05.17.100248

**Authors:** Ekin Ucuncu, Karthyayani Rajamani, Miranda S.C. Wilson, Daniel Medina-Cano, Nami Altin, Pierre David, Giulia Barcia, Nathalie Lefort, Marie-Thérèse Vasilache-Dangles, Gaële Pitelet, Elsa Lorino, Nathalie Rabasse, Eric Bieth, Maha S. Zaki, Meral Topcu, Fatma Mujgan Sonmez, Damir Musaev, Valentina Stanley, Christine Bole-Feysot, Patrick Nitschké, Arnold Munnich, Nadia Bahi-Buisson, Catherine Fossoud, Fabienne Giuliano, Laurence Colleaux, Lydie Burglen, Joseph G. Gleeson, Nathalie Boddaert, Adolfo Saiardi, Vincent Cantagrel

**Affiliations:** Paris Descartes–Sorbonne Paris Cité University, Imagine Institute, 75015 Paris, France; Developmental Brain Disorders Laboratory, Imagine Institute, INSERM UMR 1163, 75015 Paris, France; MRC Laboratory for Molecular Cell Biology, University College London, WC1E 6BT London, UK; Transgenesis Platform, Laboratoire d’Expérimentation Animale et Transgenèse (LEAT), Imagine Institute, Structure Fédérative de Recherche Necker INSERM US24/CNRS UMS3633, 75015 Paris, France; Department of Genetics, Necker Enfants Malades University Hospital, APHP, 75015, Paris, France; iPSC Core Facility, Imagine Institute, INSERM UMR 1163, 75015 Paris, France; Neurologie pédiatrique, Université Paris Descartes, Hôpital Necker-Enfants Malades, 75015 Paris, France; Service de Neuropédiatrie, CHU Nice, 06200 Nice, France; ESEAN, 44200 Nantes, Service de maladies chroniques de l’enfant, CHU Nantes, 44093 Nantes, France; Service de pédiatrie, hôpital d’Antibes-Juan-les-Pins, 06600 Antibes-Juan-les-Pins, France; Service de Génétique Médicale, CHU Toulouse, 31059 Toulouse, France; Clinical Genetics Department, Human Genetics and Genome Research Division, National Research Centre, Cairo 12311, Egypt; Hacettepe University, Faculty of Medicine, Department of Child Neurology, Ankara, 06100, Turkey; Karadeniz Technical University, Faculty of Medicine, Department of Child Neurology, Trabzon, 61080, Turkey; Laboratory for Pediatric Brain Diseases, Rady Children’s Institute for Genomic Medicine, University of California San Diego, La Jolla (CA), 92093, USA; Genomics Platform, Imagine Institute, INSERM UMR 1163, 75015 Paris, France; Bioinformatics Core Facility, Imagine Institute, INSERM UMR 1163, 75015 Paris, France; Translational Genetics Laboratory, Imagine Institute, INSERM UMR 1163, 75015 Paris, France; Genetics and Development of the Cerebral Cortex, Imagine Institute, INSERM UMR 1163, 75015 Paris, France; Centre de Référence des Troubles des Apprentissages, Hôpitaux Pédiatriques de Nice CHU-Lenval, 06200 Nice, France; Service de Génétique Médicale, Centre Hospitalier Universitaire de Nice, 06202 Nice, France; Centre de Référence des Malformations et Maladies Congénitales du Cervelet, Département de Génétique, AP-HP, Sorbonne Université, Hôpital Trousseau, 75012, Paris, France; Department of Pediatric Radiology, INSERM UMR 1163 and INSERM U1000, Necker Enfants Malades University Hospital, APHP, 75015 Paris, France

## Abstract

Inositol polyphosphates are vital metabolic and secondary messengers, involved in diverse cellular functions. Therefore, tight regulation of inositol polyphosphate metabolism is essential for proper cell physiology. Here, we describe an early-onset neurodegenerative syndrome caused by loss-of-function mutations in the *multiple inositol polyphosphate phosphatase 1* gene (*MINPP1*). Patients were found to have a distinct type of Pontocerebellar Hypoplasia with typical basal ganglia involvement on neuroimaging. We found that patient-derived and genome edited *MINPP1*^-/-^ induced pluripotent stem cells (iPSCs) are not able to differentiate efficiently into neurons. MINPP1 deficiency results in an intracellular imbalance of the inositol polyphosphate metabolism. This metabolic defect is characterized by an accumulation of highly phosphorylated inositols, mostly inositol hexakiphosphate (IP_6_), detected in HEK293, fibroblasts, iPSCs and differentiating neurons lacking MINPP1. In mutant cells, higher IP_6_ level is expected to be associated with an increased chelation of intracellular cations, such as iron or calcium, resulting in decreased levels of available ions. These data suggest the involvement of IP_6_-mediated chelation on Pontocerebellar Hypoplasia disease pathology and thereby highlight the critical role of MINPP1 in the regulation of human brain development and homeostasis.

## INTRODUCTION

Inositol polyphosphate (IPs) comprise an ubiquitous family of small molecule messengers controlling every aspect of cell physiology ^1^. The most characterized is the calcium release factor inositol trisphosphate (I(1,4,5)P^3^ or simply IP^3^), a classical example of second messenger ^2^, generated after receptor activation by the action of phospholipase C on the lipid phosphoinositide PIP^2^. Each of the six hydroxyl groups of the inositol ring can be phosphorylated, and the combination of these phosphorylations generates multiple derivatives ^3^. Among them, inositol hexakisphosphate (IP_6_, or phytic acid) is the most abundant in nature. In plants, IP_6_ accumulates in seeds, within storage vacuoles, where it could represent 1-2 % of their dry weight ^4^. Plant seed IP_6_ is used as the main source of phosphate and mineral nutrients (e.g. Ca^2+^, K^+^, Fe^2+^) during germination. In mammalian cells, IP_6_ is the most abundant inositol polyphosphate species, reaching cellular concentrations of ∼15-100 μM ^5^. IP_6_ is synthesized from inositol monophosphate (IP) or from IP_3_ by the action of several inositol phosphate kinases: IPMK (Inositol Polyphosphate Multikinase, also known as IPK2), IP_3_-3K (Inositol -1,4,5-trisphophate 3-Kinase), ITPK1 (Inositol Tetrakisphosphate 1-Kinase) and IPPK (Inositol-Pentakisphosphate 2-Kinase, also known as IPK1)^1 6^. Subsequently, the fully phosphorylated ring of IP_6_ can be further phosphorylated to generate the more polar inositol pyrophosphates such as IP_7 7_. While IP_6_ anabolism is well studied, its catabolism has been less characterized. Mammalian cells dephosphorylate IP_6_ through the action of the MINPP1 (Multiple Inositol-Polyphosphate Phosphatase 1) enzyme ^8^ that is able to degrade IP_6_ to IP_3 9_. The analysis of mouse knockouts for the inositol kinases responsible for IP_6_ synthesis have highlighted an important role for this pathway in controlling central nervous system development, since knockout of *Itpk1* or *Ipmk* is embryonically lethal due to improper neural tube development ^10 11^. In mammals, IP_6_ has been directly associated with a pleiotropy of functions, including ion channel regulation, control of mRNA export, DNA repair, and membrane dynamics ^1^. Furthermore, IP_6_ is considered as a natural antioxidant since its iron-chelating property enables it to inhibit iron-catalyzed radical formation^12^. Although not yet thoroughly studied, some of the physiological roles of IP_6_ could be related to its high affinity for polyvalent cations ^13 14^. To investigate the role of IP_6_ in mammalian physiology, many studies use IP_6_ exogenously added to cell lines in culture, often observing antiproliferative properties ^15^. These studies give little attention to the chelating property of IP_6_: cations-IP_6_ precipitation depletes the medium of essential ions such as calcium or iron. Additionally, the physiological relevance of extracellular IP_6_ in mammals is not established. Extracellular pools of IP_6_ have only been demonstrated in a cestode intestinal parasite ^16^, and several studies suggest that dietary IP_6_ cannot be absorbed as such through the digestive system and is absent from body fluids ^17 18^. Instead, *de novo* synthesis of IP_6_ occurs in all mammalian cells, including in the brain with high levels in regions such as the brainstem and striatum ^17 19^. The existence of several cellular pools of IP6 has been suggested ^19 20 6^. However, the dynamic regulation of the endogenous intracellular pools of IP_6_ is not fully understood, since its high cellular concentration precludes the determination of IP_6_ pool specific fluctuations. Therefore, the exact function(s) of IP_6_ in cell homeostasis and mammalian development remain an area of intense investigation.

Several human diseases have been genetically associated with alterations in phosphoinositide (the lipid derivatives of inositol) metabolism ^21^. However, so far, no Mendelian disorder has been shown to be caused by an imbalance in the cytosolic inositol polyphosphate pathway, with the exception of a single variant in a gene involved in the conversion of the pyrophosphates forms of inositol and associated with hearing impairment ^22^.

Pontocerebellar hypoplasia (PCH) is a group of early-onset neurodegenerative disorders that includes at least 13 subtypes, based on neuropathological, clinical and MRI criteria ^23 24^. PCH is usually associated with a combination of degeneration and lack of development of the pons and the cerebellum, suggesting a prenatal onset. The genetic basis is not known for all of the cases, and preliminary data from different PCH cohorts suggest that many subtypes remain to be identified. Based on the known molecular causes, PCH often results from a defect in apparently ubiquitous cellular processes such as RNA metabolism regulation and especially tRNA synthesis (i.e. mutations in *EXOSC3, TSEN54, TSEN2, TSEN34, CLP1* and *RARS2*). Multiple additional conditions show neurological symptoms and imaging comparable to typical PCH syndromes and are caused by defects in diverse pathways involved in mitochondrial, glycosylation or purine nucleotide metabolisms. This observation further supports the disruption of ubiquitous pathways as the unexplained basis of these neurological conditions^23^.

In this study, we identify MINPP1 as necessary for the dephosphorylation of intracellular IP_6_, and describe a new syndrome of PCH caused by a defect in this process that directly regulates cytosolic cation (e.g. Ca^2+^, Fe^3+^) homeostasis.

## RESULTS

### Loss of function mutations of the *MINPP1* gene are associated with a distinct subtype of Pontocerebellar Hypoplasia

To identify new etiological diagnoses of patients with PCH, we explored a group of 15 probands previously screened negative with a custom gene panel approach ^25^. Whole exome sequencing (WES) was then performed through trio sequencing (i.e. both parents and the proband). Among the new candidate genes that were identified, the *MINPP1* gene was recurrent and the most obvious candidate (Table 1 and Supplementary Note). The *MINPP1* gene has not been previously associated with any Mendelian disorders. To assess how frequently *MINPP1* mutations could be involved in PCH, we explored two other cohorts of pediatric cases with neurological disorders. The presence of *MINPP1* mutations was investigated using a custom gene panel or WES. Three additional families with *MINPP1* bi-allelic variants were identified, all the affected being diagnosed with PCH.

**TABLE 1:**
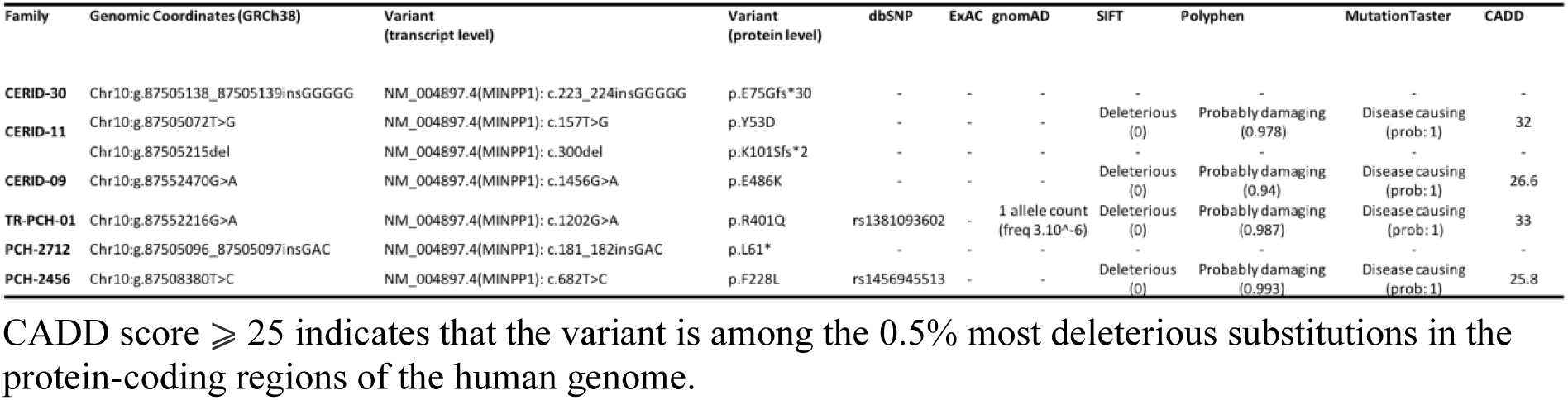
*MINPP1* variants identified in PCH cases.

| Family | Genomic Coordinates (GRCh38) | Variant (transcript level) | Variant (protein level) | dbSNP | ExAC | gnomAD | SIFT | Polyphen | MutationTaster | CADD |
| --- | --- | --- | --- | --- | --- | --- | --- | --- | --- | --- |
| CERID-30 | Chr10:g.87505138_87505139insGGGGG | NM_004897.4(MINPP1): c.223_224insGGGGG | p.E75Gfs*30 | - | - | - | - | - | - | - |
| CERID-11 | Chr10:g.87505072T>G | NM_004897.4(MINPP1): c.157T>G | p.Y53D | - | - | - | Deleterious (0) | Probably damaging (0.978) | Disease causing (prob: 1) | 32 |
|  | Chr10:g.87505215del | NM_004897.4(MINPP1): c.300del | p.K101Sfs*2 | - | - | - | - | - | - | - |
| CERID-09 | Chr10:g.87552470G>A | NM_004897.4(MINPP1): c.1456G>A | p.E486K | - | - | - | Deleterious (0) | Probably damaging (0.94) | Disease causing (prob: 1) | 26.6 |
| TR-PCH-01 | Chr10:g.87552216G>A | NM_004897.4(MINPP1): c.1202G>A | p.R401Q | rs1381093602 | - | 1 allele count (freq 3.10^-6) | Deleterious (0) | Probably damaging (0.987) | Disease causing (prob: 1) | 33 |
| PCH-2712 | Chr10:g.87505096_87505097insGAC | NM_004897.4(MINPP1): c.181_182insGAC | p.L61* | - | - | - | - | - | - | - |
| PCH-2456 | Chr10:g.87508380T>C | NM_004897.4(MINPP1): c.682T>C | p.F228L | rs1456945513 | - | - | Deleterious (0) | Probably damaging (0.993) | Disease causing (prob: 1) | 25.8 |

CADD score $\geq 25$ indicates that the variant is among the 0.5% most deleterious substitutions in the protein-coding regions of the human genome.

In total, bi-allelic variants in *MINPP1* were identified in eight affected children from six unrelated families (Fig.1, Table 1, Supplementary Fig.1). These variants include homozygous early-truncating mutations in the families CerID-30 and PCH-2712, compound heterozygous missense and frameshift variants in family CerID-11, a homozygous missense variant in the endoplasmic reticulum (ER) retention domain of the protein in the family CerID-09 and homozygous missense variants in the histidine phosphatase domain of the protein in the families TR-PCH-01 and PCH-2456 (Fig.1B, D). These four missense variants are predicted to be disease-causing using MutationTaster and SIFT ^26^, and involve amino acids fully conserved across evolution (Table 1, Fig.1C). To predict the impact of the variants on protein structure, we utilized a crystal structure of D. castellii phytase and evaluated the consequences of the missense variants involving amino-acids included in the model (Supplementary Fig.1B). Tyr53Asp variant introduces a buried charge and disrupts a hydrogen bond with the donor amino-acid Ser299. The Phe228Leu substitution breaks a buried hydrogen bond with Lys241, both amino-acid positions are close to the IP_6_ binding site. The Arg401Gln substitution replaces a buried charged residue with an uncharged residue and disrupts a salt bridge formed with the amino-acid Asp318. Thus, all the missense variants tested are predicted to cause structural damages with potential consequences on the enzyme activity.

**Figure 1:**
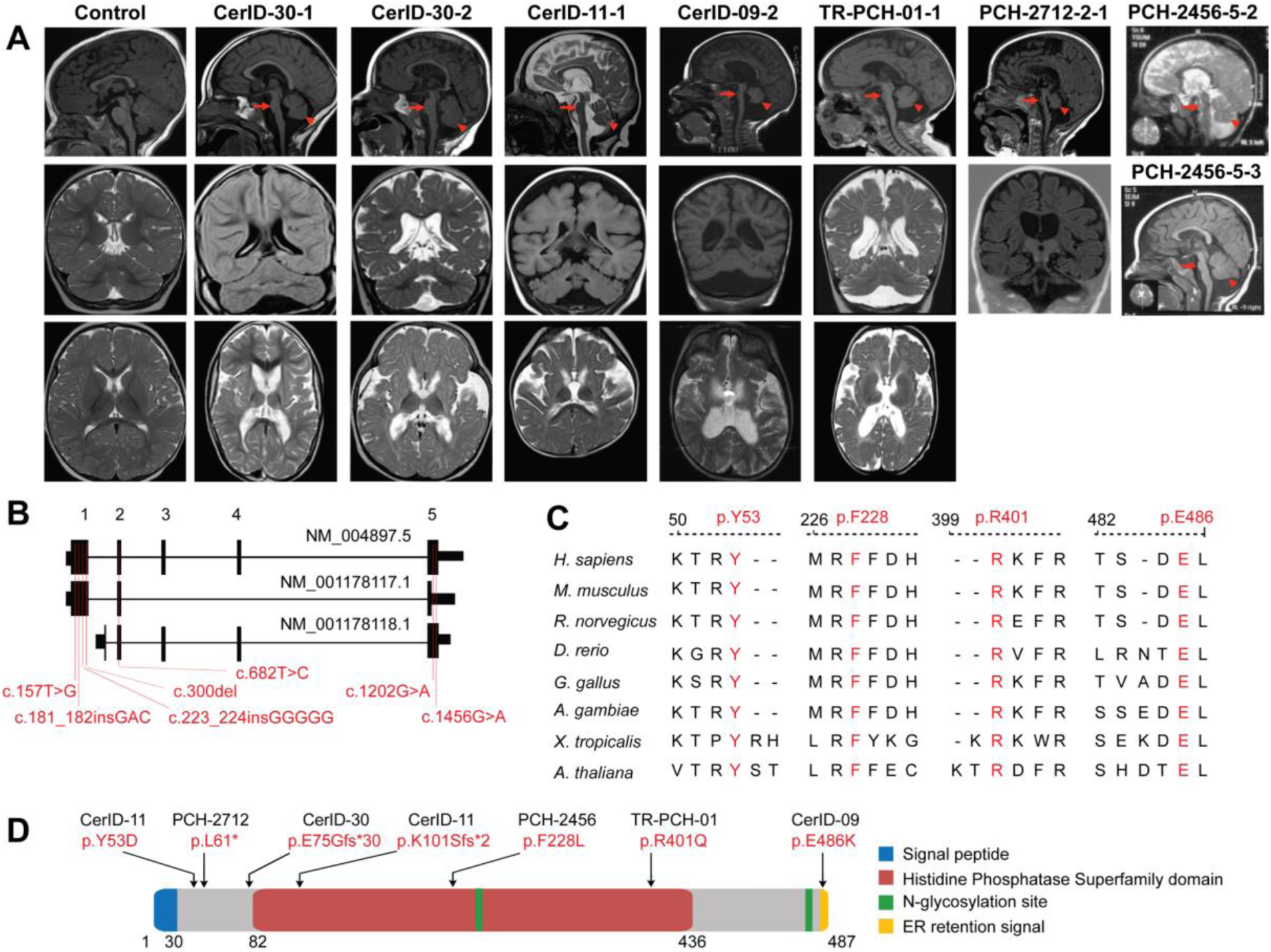
Biallelic mutations in *MINPP1* cause a distinct PCH phenotype. **(A)** Midline sagittal (top), coronal (middle) and axial (bottom) brain MRIs of control and patients from families CerID-30, CerID-11, CerID-09 and TR-PCH-01 respectively. Only sagittal (top) and coronal (middle) brain MRIs were available for the patient from the family PCH-2712 and sagittal brain MRI for the patients from PCH-2456 (top and middle). Sagittal MRIs show variable degree of brain stem (arrow) and cerebellar atrophy/hypoplasia (arrowhead). (**B)** Schematic representation of the *MINPP1* transcripts: NM_004897.5, NM_001178117.1 and NM_001178118.1 respectively. Exon numbers for the longest isoform NM_004897.5 are indicated above the schematic representation. Mutations are shown relative to their cDNA (NM_004897.5) position. (**C)** Multiple-sequence alignment of MINPP1 from different species. Variant amino-acid residues p.Y53, p.F228, p.R401 and p.E486 are evolutionarily conserved. (**D)** Linear schematic representation of MINPP1, showing the position of mutations with respect to predicted protein domains. Abbreviations used: Endoplasmic reticulum (ER).

The eight patients presented with almost complete absence of motor and cognitive development, progressive or congenital microcephaly, spastic tetraplegia or dystonia, and vision impairments (Table 2). For most of the patients, the first symptoms included neonatal severe axial hypotonia and epilepsy that started during the first months or years of life. Pre-natal symptoms of microcephaly associated with increased thalami echogenicity were detected for the individual CerID-11, while the seven other patients presented with progressive microcephaly. For patients from the families CerID-09 and 11, the phenotype appeared to be progressive and the affected children died in their infancy or middle-childhood. Strikingly, all the affected children harbor a unique brain MRI showing a mild to severe PCH, fluid-filled posterior fossa, with dilated lateral ventricles. Additionally, severe atrophy at the level of the basal ganglia or thalami often associated with typical T2 hypersignal were identified in all the patients MRI (Table 2, Fig.1A lower panel). This imaging is distinct from other PCH syndromes and thereby defines a new subtype that we propose as PCH type 14.

**TABLE 2:**
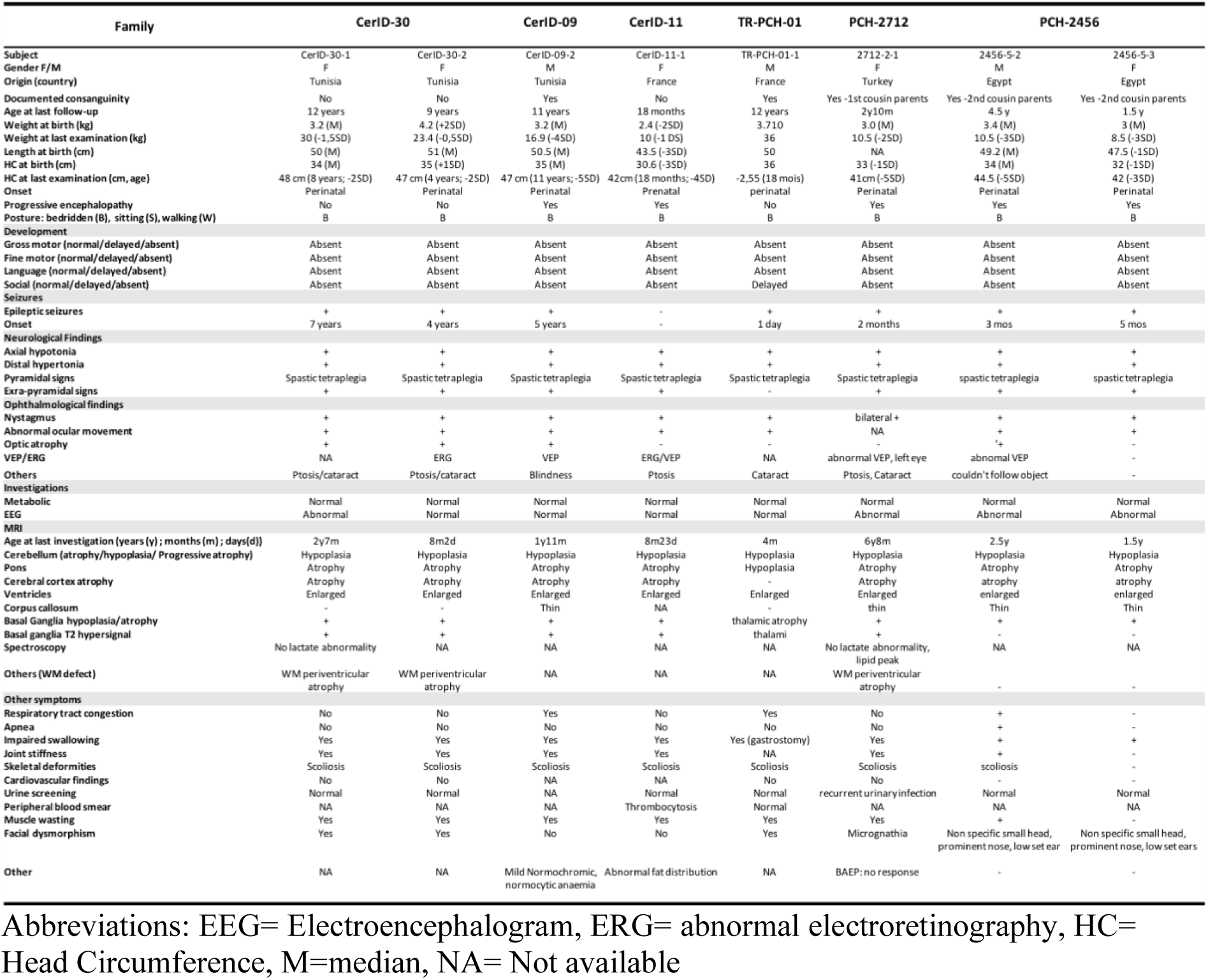
Clinical information of the patients with variants in *MINPP1*.

### *MINPP1* missense PCH mutations are deleterious for protein function

The MINPP1 enzyme is predominantly localized in the ER lumen^27^. It removes phosphate groups from inositol polyphosphate substrates starting at position 3 ^28,29^, with high affinity for IP_5_ and IP_6 30_. Indeed, it has been described as the main mammalian phytase, or enzyme involved in IP_6_ degradation (Fig. 2A). Despite its name, this gene does not have any paralog in the human or mouse genome. In order to determine the effect of the patient mutations on the endogenous enzyme, we obtained skin fibroblast from patients of the CerID-30 family. MINPP1 protein was undetectable in patients’ cells (Fig. 2B), supporting a complete loss of function of *MINPP1* as the cause of this PCH subtype. The *MINPP1* gene is widely expressed in the developing and adult mouse ^8 30^, and rat^9^, as well as in human tissues (Supplementary Fig. 2A). This broad expression pattern suggests a general role for this enzyme in regulating inositol polyphosphate metabolism. To investigate this role, we generated a HEK293T cell KO model for the *MINPP1* gene (*MINPP1*^-/-^ HEK293) using genome editing (Fig. 2C). *MINPP1*^-/-^ HEK293 cells showed a 30% decrease in their growth rate after 48 hours of culture, likely resulting from a proliferation defect, in the absence of significant difference in the binding of the apoptosis marker Annexin-V (Fig.2D, Supplementary Fig.2B, C). This defect was partially rescued by transient over-expression of WT MINPP1 (p<0.001; Fig. 2E, Supplementary Fig.2D). Contrastingly, over-expression of two of the MINPP1 missense variants (i.e. Y53D and E486K) did not rescue growth, suggesting that these variants have a major impact on the protein function. MINPP1 has two predicted N-glycosylation sites (Fig.1D). In order to evaluate the glycosylation status of the endogenous and over-expressed WT MINPP1 as well as the Y53D and E486K missense variants, we treated the protein extracts with the PNGase enzyme ^31^. A shift in the molecular weight after treatment, in all the samples, indicated that MINPP1 is indeed N-glycosylated. No major changes were observed in presence of the missense variants although a decrease in the amount of the fully glycosylated MINPP1 with the E486K variant can not be excluded (Supplementary Fig.2E).

**Figure 2:**
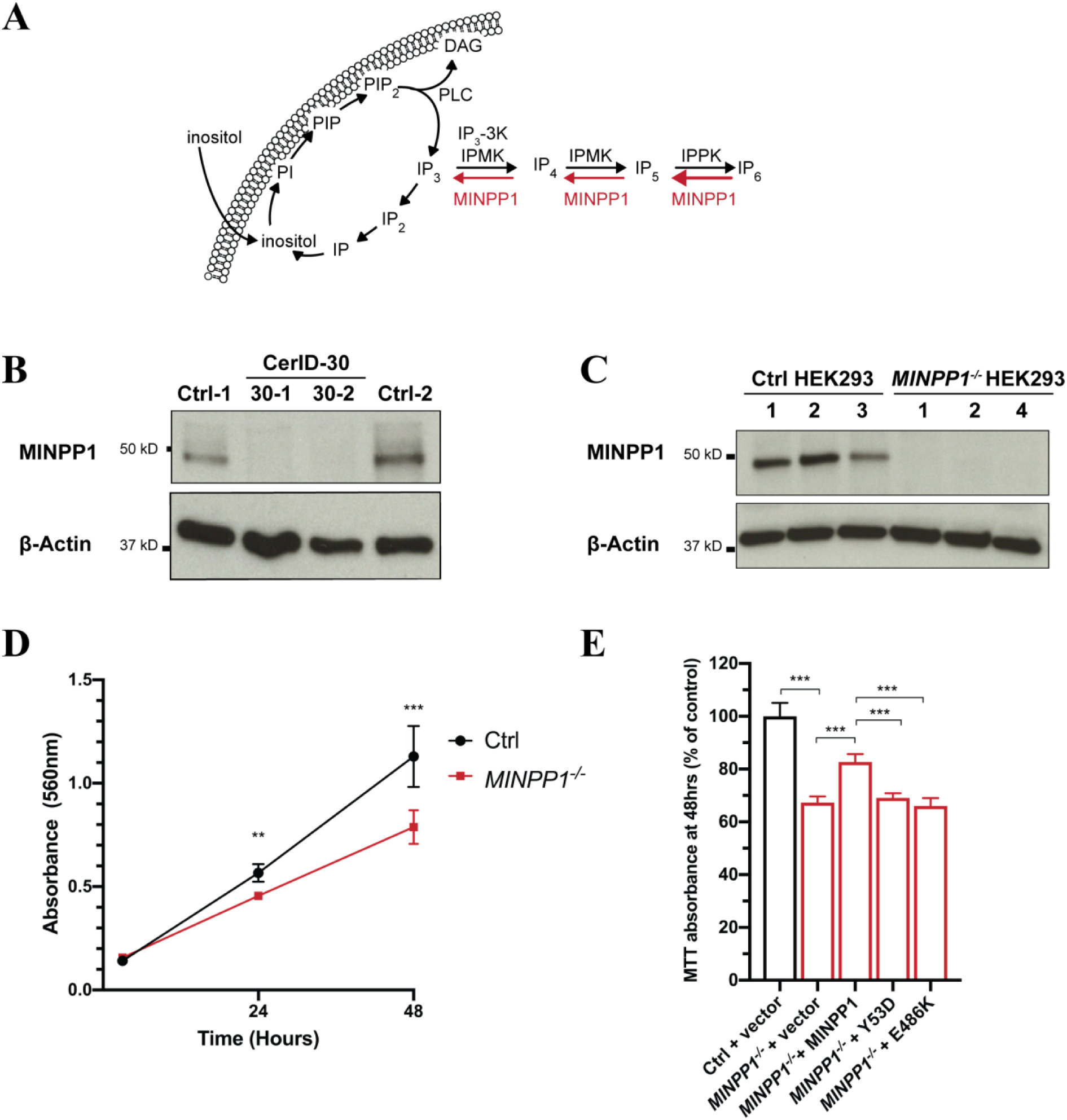
PCH-associated mutations of *MINPP1* are deleterious for protein function. **(A)** Schematic representation of inositol phosphate cycle. Abbreviations used: *myo-*inositol (Inositol) phosphatidylinositol (PI); phosphatidylinositol phosphate (PIP); phosphatidylinositol 4,5-bisphosphate (PIP_2_); diacylglycerol (DAG); Phospholipase C (PLC); inositol phosphate (IP); inositol bisphosphate (IP_2_); inositol 1,4,5-trisphosphate (IP_3_); inositol 1,3,4,5-tetrakisphosphate (IP_4_); inositol pentakisphosphate (IP_5_); inositol hexakisphosphate (IP_6_); Inositol Polyphosphate Multikinase (IPMK); I(1,4,5)P_3_ 3-Kinase (IP_3_-3K); Inositol-Pentakisphosphate 2-Kinase (IPPK); Multiple Inositol-Polyphosphate Phosphatase 1 (MINPP1). (**B-C**) Western blot analysis of MINPP1 level in patient fibroblasts and HEK293 cells with β-Actin shown as loading control. Patient fibroblasts CerID-30-1 and CerID-30-2 (**B**) and *MINPP1*^*-/-*^ HEK293 clones (**C**) show absent MINPP1. (**D)** Assessment of cell proliferation by MTT assay. For each clone, MTT absorbance was measured 3, 24 and 48 hours post-seeding. Values represent the mean ± s.d. of triplicate determinations from 4 replicates. (n=4, Two-tailed student’s t-test, ** and *** indicate the p values p<0.01 and P ≤ 0.001 respectively). **(E)** *MINPP1*^*-/-*^ HEK293 cells were transiently transfected with plasmids encoding empty vector, wildtype, Y53D or E486K variant MINPP1. To assess the cell proliferation rate, MTT assay was performed 48 hours post-nucleofection. The data are presented as mean percentage relative to control (Ctrl) ± s.d., with triplicate determinations from four replicates. The normalization was done with 3 hours MTT assay data. (n=4, One-way ANOVA, Tukey’s post hoc test, *** indicates p value p≤0.001).

### Patient and genome-edited *MINPP1* mutant iPSCs show an impaired neuronal differentiation

To investigate the mechanism at the origin of the neurological symptoms of *MINPP1* patients, we derived induced pluripotent stem cells (iPSCs) from patient CerID-30-2 (Supplementary Fig.3A, B). In order to assess a contribution of the genetic background or other factors to the phenotype, we also generated *MINPP1*^-/-^ iPSCs in isogenic background (Supplementary Fig. 3A, B). Surprisingly, a dual SMAD inhibition-based neural induction protocol^32 33^, did not allow the generation of viable neural progenitor cells for both *MINPP1* mutant lines (data not shown). Differentiation of patient-derived iPSCs systematically generated mixed cell populations with undefined HNK1 negative cells (Supplementary Fig. 3C). These observations suggest a critical role for MINPP1 during neuroectodermal induction, and led us to use a different protocol that preserved neural rosette environment, using only noggin as a SMAD inhibitor ^34^. In these conditions, control cells efficiently differentiated toward TUJ1^+^ neurons after 14 days (Fig. 3A). In contrast, both *MINPP1* mutant lines showed significant 42% and 65% decreases in TUJ1^+^ post-mitotic cells mirrored by significant 2.6 and 3.1-fold increases in the number of PAX6^+^ neural progenitors in *MINPP1*^-/-^ and CerID-30-2 derived cells respectively (Fig. 3A, B). These changes reflect the inability of neural progenitors to efficiently differentiate into post-mitotic neurons. Altogether, these observations show that *MINPP1* plays a direct role during human neuronal differentiation, and suggest that a differentiation defect could be involved in the neuronal vulnerability underlying this early-onset neurodegenerative disorder.

**Figure 3:**
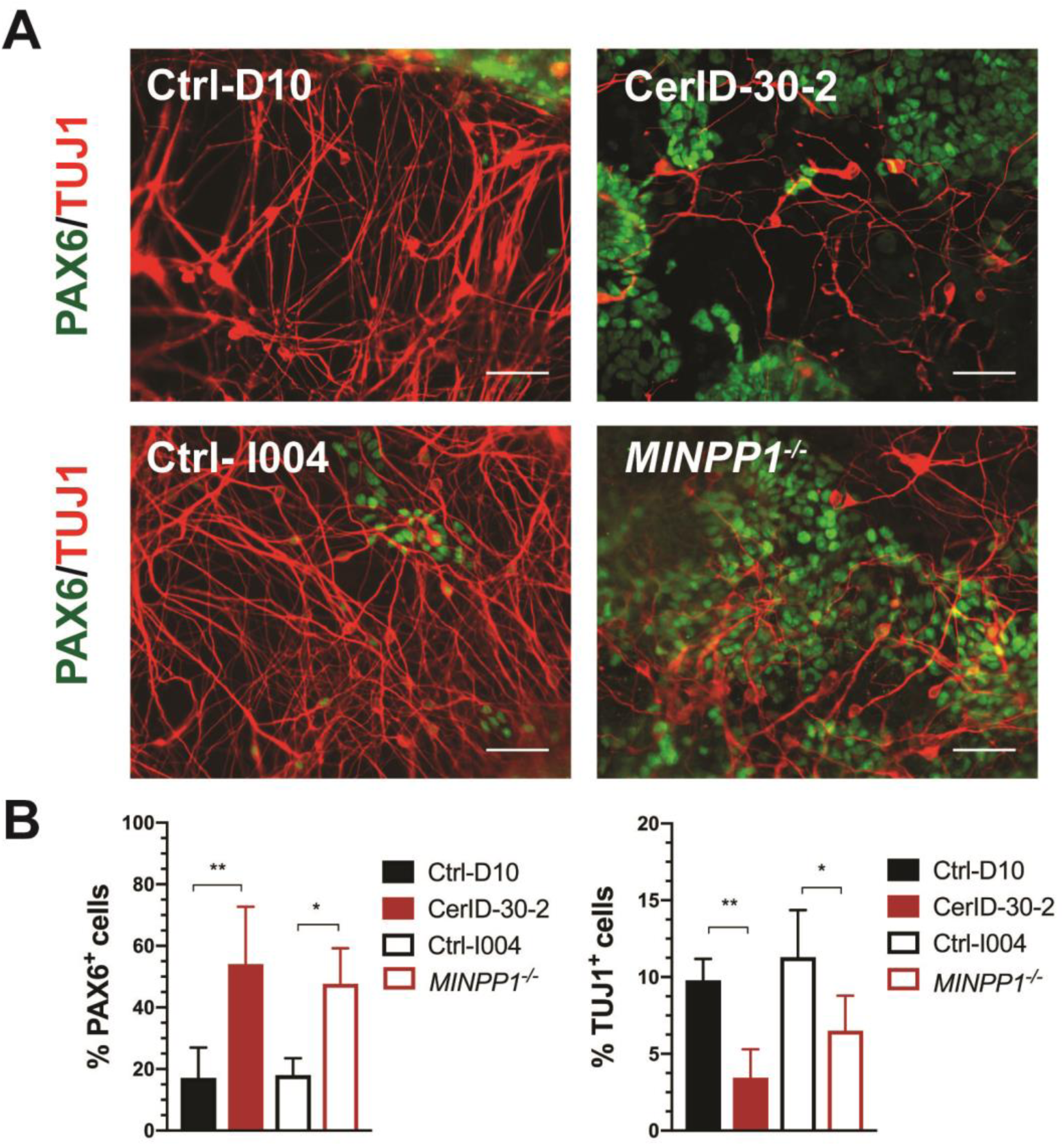
MINPP1 absence impairs early neuronal differentiation. **(A)** Control (Ctrl-D10 and Ctrl-I004), patient-derived (CerID-30-2) and *MINPP1*^-/-^ iPSCs were differentiated towards neuronal lineage for 14 days. Representative images of the differentiated cells stained with early neuronal marker TUJ1 and neural progenitor marker PAX6. Hoechst was used as a nuclear stain. All scale bars correspond to 50μm. **(B)** Quantitative analysis of the immunofluorescence data. (Duplicate analysis of two independent experiments, one-way ANOVA, Tukey’s post hoc test, * and ** indicate p value: p<0.05 and p<0.01). The data are presented as mean percentage values ± s.d.

### Inositol polyphosphate metabolism is altered in HEK293, iPSCs and induced neurons mutated for *MINPP1*

The localization of human MINPP1 into the ER, and the demonstration that its drosophila homolog (i.e. mipp1) is anchored to the plasma membrane outside of the cell^35^, prompted us to investigate the presence of phytase activity in conditioned media from control and *MINPP1* mutant HEK293 cells. In conditioned medium from control cells, exogenously added IP_6_ was substantially processed after two hours, and completely degraded after four hours (Supplementary Fig.4A). Conversely, although partially degraded, IP_6_ is still detectable after six hours of incubation in *MINPP1* mutant conditioned media. This result suggests that MINPP1 accounts for the main secreted phytase activity of HEK293 cells.

To explore precisely a disruption in inositol phosphate metabolism, and to better address the role of MINPP1 in this metabolic pathway, we used tritium inositol (*myo*-[^3^H]-inositol) metabolic labeling of cultured cells, and analyzed inositol derivatives with SAX-HPLC (strong anion-exchange high-performance liquid chromatography) as previously described ^36 37^. We applied this method to HEK293, skin fibroblasts, and iPSCs before or during neuronal differentiation at day 10 (referred to as Day 10 differentiating neurons) from control and *MINPP1* mutant cell lines. Exogenously added [^3^H]-inositol is imported into the cytosol and converted into phosphoinositide lipids before processing into inositol phosphates (IPs) (Fig. 2A). As expected, after 3 days of [^3^H]-inositol labeling, IP_6_ was detected as the most, or the second most abundant intracellular inositol derivative in control cell lines (hollow and filled bars in Fig.4), but was absent in the cell culture media (Fig.4, Supplementary Fig.4B). In all the cell models studied, the disruption of MINPP1 enzyme activity had a strong impact on intracellular IPs profile when compared with their respective controls. The investigation of *MINPP1*^-/-^ HEK293 cells revealed a 3-fold significant increase in IP_6_ level, as well as an increase in IP_5_ levels, and surprisingly a severe decrease in IP and IP_2_ levels (Fig.4A). Trends in the same direction, although not significant, were detected in patient fibroblasts (Fig.4B) suggesting cell-type specific differences. Indeed, in iPSCs, IP_6_ levels showed a significant 1.6-fold increase in both patient-derived and *MINPP1* KO iPSCs, also associated with an increase in IP_5_ levels (Fig.4C, D). Finally, the study of Day 10 differentiating neurons revealed significant 1.9-fold and 1.6-fold increases in IP_6_ levels in patient-derived and *MINPP1* KO differentiating cells respectively, with a trend or significant decrease in IP_2_ levels (Fig.4E, F). Interestingly, the defects in IP_6_ and IP_2_ levels in *MINPP1*^*-/-*^ HEK293 cells were fully rescued, considering a ∼70% transfection efficiency, by the transient over-expression of WT MINPP1 (Supplementary Fig. 4C and D). We also performed radiolabeling experiments on *MINPP1*^*-/-*^ HEK cells with Y53D and E486K MINPP1 variants. The Y53D variant had no impact on the IP_6_ and IP_2_ levels, indicating the clear loss of enzyme activity (Supplementary Fig.4D). However, the E486K variant did not affect the enzyme activity in this overexpression system, with apparently unchanged levels of IP_6_ and IP_2_. This result is in line with the previously demonstrated preserved enzyme activity in the absence of the ER retention peptide ^30^. Therefore, and given its inability to rescue the *MINPP*1^-/-^ HEK293 proliferation, this variant could impact a critical regulation of MINPP1 protein.

**Figure 4:**
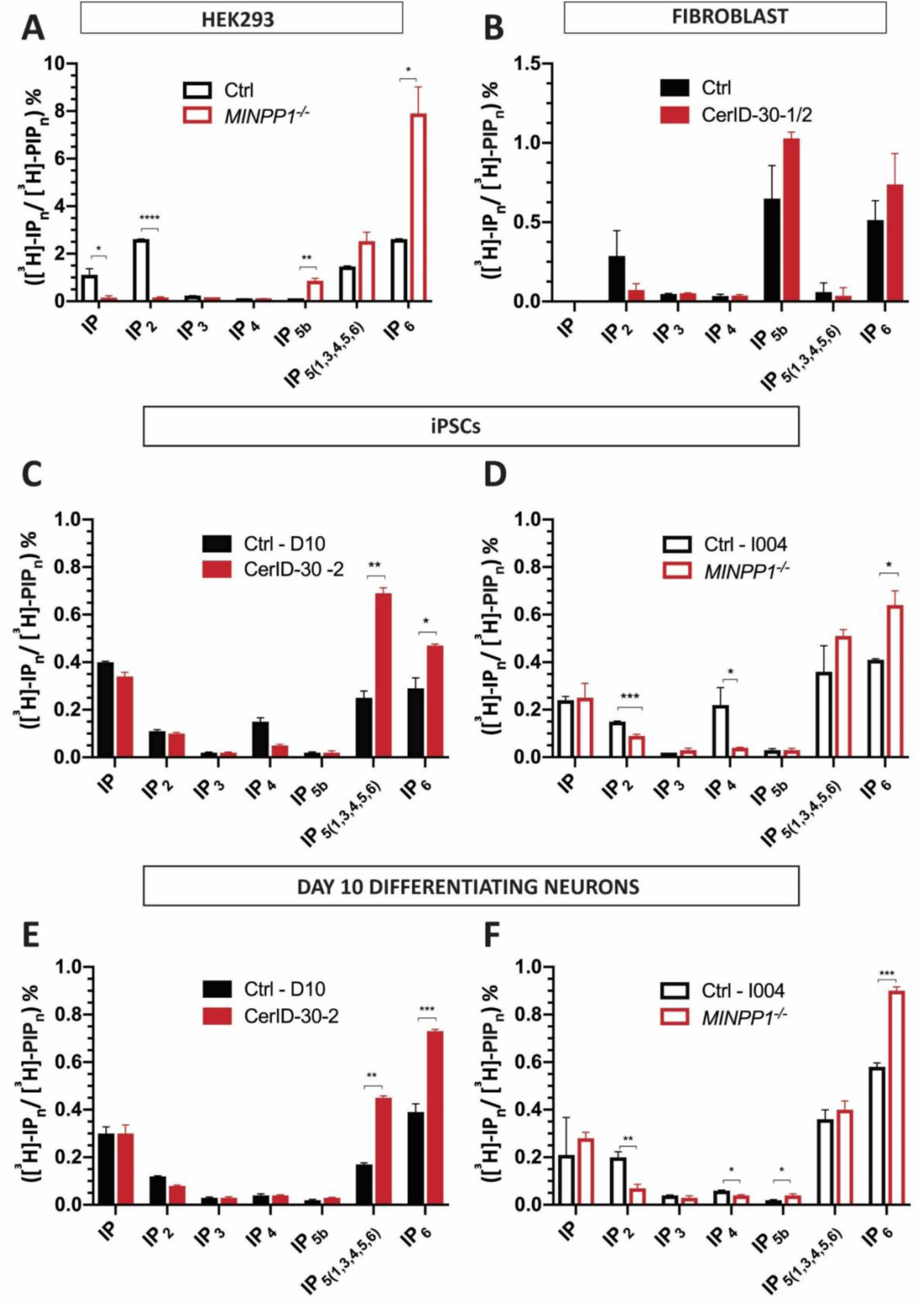
MINPP1 absence leads to disruption in inositol phosphates metabolism. **(A-F)** SAX-HPLC analysis of inositol phosphate levels in *MINPP1*^-/-^ HEK293 cells **(A)**, patient fibroblasts (CerID-30-1 and CerID-30-2) **(B)**, patient-derived (CerID-30-2) **(C)** and *MINPP1*^-/-^ iPSCs **(D)**, and their day-10 differentiating neuron counterparts **(E-F)**. The peaks ([^3^H]-IP_n_) were identified based on comparison to standards. [^3^H]-IP_n_ levels are presented as percentage of total radioactivity in the inositol-lipid fraction ([^3^H]-PIP^n^). All error bars represent standard deviation (s.d.). (n=2, Two-tailed student’s t-test for HEK293 and fibroblast data; One-way ANOVA for iPSCs and their differentiated counterparts data, Tukey’s post hoc test, *, **, ***, **** indicate p values p<0.05, p<0.01, p<0.001, p ≤ 0.0001 respectively). Abbreviations used: IPn, inositol phosphates; PIPn, phosphatidyl inositol phosphates.

In all the cell models tested, including HEK293 cells, patients’ fibroblasts, and undifferentiated and differentiated iPSCs, a comparable imbalance of IPs levels were observed, where increase in the amounts of higher inositol polyphosphate derivatives IP_5_ and IP_6_ were associated with a decrease in lower-phosphorylated IP_2_ and IP species (Fig. 4). Differences observed between the various cell models tested are likely to be caused by cell type differences and potentially also by the genetic background. Nevertheless, these observations clearly demonstrate the critical role played by MINPP1 in cellular inositol polyphosphate homeostasis, with the conversion of higher to lower IPs. The most robust finding was that IP_6_ is systematically increased in *MINPP1* mutant cells compared to controls. Altogether, these observations exclude a major contribution of extracellular higher-phosphorylated IPs to this metabolic defect, but highlight an unappreciated role for *MINPP1* in the regulation of the intracellular pool of de novo synthesized IP_6_, the most abundant inositol derivative.

### IP_6_ accumulation can deplete free iron in presence of high iron condition

Considering the strong impact of *MINPP1* mutations on cellular IP_6_ levels, and the known chelator properties of this molecule, we hypothesized that MINPP1 defects can have consequences for intracellular cations homeostasis. An intracellular accumulation of IP_6_ could theoretically lead to the accumulation of chelated cations inside the cell, potentially reducing the pool of free cations. At physiological pH, IP_6_ has a strong binding affinity to iron^14^, therefore we evaluated the ability of HEK293 cells to store iron, in low iron (-FAC) or high iron (provided with ferric ammonium citrate; +FAC) conditions. Then, we used a colorimetric ferrozine-based assay with a HCl/KMnO_4_ pretreatment step that separates iron from its binding molecules to measure total intracellular iron ^38 39^. After two days of incubation with FAC, we observed a significant 1.5-fold increase in the total iron content in *MINPP1*^-/-^ HEK293, under high iron conditions compared to control (Fig. 5A). Although based on non-physiological iron conditions, this observation suggests that IP_6_ could play a role in the regulation of metal ion cellular storage such as iron. To investigate a potential increase of iron chelation affecting the free iron cellular pool, we measured the cellular free Fe^2+/3+^ content using standard cell lysis and colorimetric assay. We detected a 58% depletion in total free iron levels in *MINPP1* mutant cells under high iron conditions (Fig.5B). Interestingly, this depletion was mainly contributed by a decrease in Fe^3+^ levels (Fig.5C, D). IP_6_, the major IPs accumulating in mutant HEK293, is known to have higher affinity for Fe^3+^ versus Fe^2+ 40,41^. These data are consistent with a massive accumulation of complexed iron in the absence of the MINPP1 enzyme, in the presence of high iron, and suggest the potential involvement of IP_6_-mediated abnormal cation homeostasis as the underlying disease mechanism.

**Figure 5:**
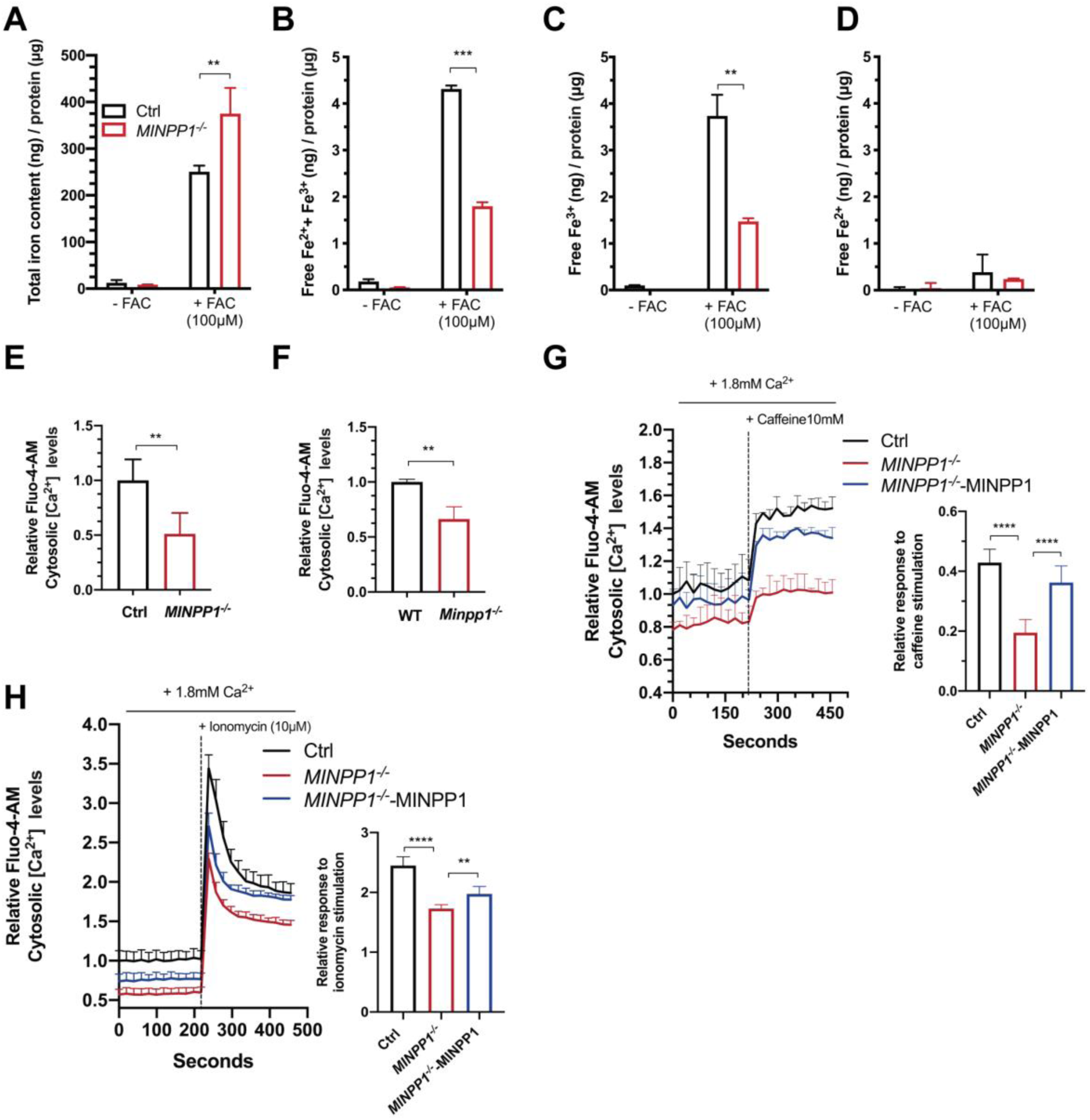
Altered iron and calcium homeostasis in the absence of MINPP1 enzyme in HEK293 and *Minpp1*^-/-^ mouse neural progenitor cells. **(A-D)** Quantification of total iron content (**A**), free iron (Fe^2+^ and Fe^3+^) (**B**), Fe^3+^ (**C**) and Fe^2+^ (**D**) levels in extracts from control and *MINPP1*^*-/-*^ HEK293 cells grown under low (-FAC) and high iron (+FAC, 100 µM) conditions. All values are normalized to the total protein concentration and represent the mean ± s.d (n=3, Two way ANOVA Sidak test, **, *** indicate p value p<0.01 and p<0.001 respectively). (**E-F**) Relative Fluo-4-AM cytosolic Ca^2+^ levels in control and *MINPP1*^*-/-*^ HEK293 cells **(E)**, wild-type (WT) and *Minpp1*^-/-^ E14 mouse neural progenitors **(F). (G-H)** Relative Fluo-4-AM cytosolic Ca^2+^ levels in control, *MINPP1*^*-/-*^ and MINPP1 over-expression stable line in *MINPP1*^-/-^ HEK293 cells (*MINPP1*^-/-^-MINPP1) loaded either with 10 mM caffeine (**G**) or 10 µM ionomycin (**H**). The dotted line indicates the addition of either caffeine (**G**) or ionomycin (**H**). Relative response after caffeine or ionomycin stimulation (peak) is represented graphically (inset). For all of the calcium assay experiments, the data are normalized to cell number with MTT colorimetric assay and presented as mean values relative to control baseline fluorescence intensity control ± s.d. ((**E**) n=5; (**F**) N=3 mice, (**G-H**) n=6, Two-tailed student’s t-test (E-F) and one-way ANOVA, Tukey’s post hoc test (G-H), ** and **** indicate p value p<0.01 and p<0.0001 respectively).

### IP_6_ accumulation causes cytosolic calcium depletion in mutant HEK293 and primary mouse neural progenitors

To further explore the involvement of the disruption of cellular cations homeostasis in PCH, we evaluated the intracellular free Ca^2+^ levels in the absence of MINPP1, using the FLUO-4-AM calcium binding indicator. Strikingly, we found a significant, close to 50% depletion of free basal Ca^2+^ levels in *MINPP1*^-/-^ HEK293, compared to control cell line (Fig. 5E). To further validate the involvement of such calcium depletion in the neurological phenotype, we generated a CRISPR/Cas9-mediated *Minpp1*^-/-^ mouse model (Supplementary Fig. 5A-E). *Minpp1* KO mice were fertile and are born at Mendelian ratio (data not shown) as described in a previously generated *Minpp1* KO mouse model^30^. Brain histology did not identify major differences in cerebellar (Supplementary Fig.5C) or cerebral cortex architecture (not shown). However, we identified a mild but significant ∼10% decrease in the brain weight associated with a reduced cortical thickness in homozygous mutant mice at P21 (Supplementary Fig.5D, E). This observation suggests the presence of an evolutionarily conserved requirement for MINPP1 activity in mammalian brain development. To identify a potential cause for this cortical phenotype, we isolated neural progenitors at E14.5 and measured the intracellular free Ca^2+^ levels. Surprisingly, we observed a significant 33.6% decrease in intracellular free Ca^2+^ levels in the *Minpp1*^-/-^ mouse cells when compared to wild type neural progenitors (p=0.007; Fig. 5F).

To assess the effects of IP_6_ accumulation on calcium signaling, we studied the caffeine sensitive ER calcium release in the *MINPP1*^-/-^ HEK293 cells^42^. In response to 10 mM caffeine, we observed a significant peak in the cytosolic Ca^2+^ levels within a minute in the control cells. However, we could only detect a slight increase with a sustained plateau in *MINPP1*^-/-^ HEK293 cells, indicating an altered response potentially caused by a decrease of Ca^2+^ in intracellular ER stores (Fig.5G). To further study the Ca^2+^ mobilization in *MINPP1*^-/-^ HEK293 cells, we treated the cells with ionomycin as it is known to initially increase the cytoplasmic calcium levels, which in turn activates calcium induced calcium response and eventually causes Ca^2+^ depletion in the ER^43^. In response to ionomycin, the control and *MINPP1*^-/-^ HEK293 cells exhibited an initial Ca^2+^ peak with a sustained plateau. However, the relative response to ionomycin stimulation was again significantly decreased in *MINPP1*^-/-^ cells (Fig.5H).

These calcium signaling defects were specifically rescued in a *MINPP1*^-/-^ HEK293 line with stable expression of MINPP1 (Fig.5G and H). Interestingly, we observed similar results in the absence of extracellular calcium (Supplementary Fig. 5F and G), for both caffeine and ionomycin, indicating that the defect is not due to the inhibition of calcium entry. Therefore, these results clearly suggest that MINPP1 absence affects the calcium levels in the cytosol as well as in the intracellular stores such as the ER. Altogether, these data support the critical role played by MINPP1 in the regulation of the intracellular IPs and available cations with strong implications for neural cell signaling and homeostasis.

## DISCUSSION

The direct physiological role(s) played by IP_6_ in mammals has been difficult to define, due to the technical challenges associated with its measurement, and its complex anabolism. Furthermore, the well described role of IP_6_ (phytate) and phytase activity in plants and bacteria had led to the thinking that extracellular IP_6_ degradation also supplies mammalian cells with phosphate and cations. On the contrary, we demonstrate that intracellular, not extracellular, IP_6_ influences cation homeostasis. An imbalance of IPs derivatives has not been so far directly involved in disease, and the previously investigated *Minpp1* KO mouse did not reveal any obvious phenotype ^30^. Surprisingly, we discovered that the absence of MINPP1 in humans results in a very severe early-onset neurodegenerative disorder with specific features. Patients with loss of function mutations in the *MINPP1* gene present with PCH associated with typical basal ganglia or thalami involvement identified by MRI. The prenatal onset of the phenotype is obvious for patient CerID-11, which supports a critical and early role for MINPP1 in neuronal development and survival. In agreement with this, we observed that patient-derived and genome-edited iPSCs mutant for *MINPP1* cannot be differentiated toward neural progenitors that efficiently give rise to neurons. Although the exact mechanism underlying this differentiation defect has not been identified yet, the sensitivity of human neural progenitors to the disruption of IPs levels is likely to be involved ^44^.

While a key role for MINPP1 in the regulation of IP_6_ cellular levels has been investigated before^30^, we provide the first evidence for its critical importance on cellular physiology and human development. Our analysis of IPs profiles using metabolic labelling unambiguously identified a typical imbalance resulting from MINPP1 defect. The increase in IP_5_ and IP_6_ levels is consistent with a previous mouse model study^8^, but our more complete assessment of the IPs metabolism imbalance also revealed alterations in lower phosphorylated IPs. Furthermore, the consequences of this metabolic block are associated with cell type-dependent differences in the IPs profile, such as the IP_4_ depletion in mutant iPSCs and robust IP_6_ accumulation in day-10 differentiating neurons mutated for *MINPP1*.

The discrepancy related to the supposed mostly cytosolic localization of IP_6_ and the ER localization of MINPP1 remains an unsolved problem ^19, 26, 30^. Hypothetically, the specific subcellular localization of MINPP1 prevents IP_6_ accumulation in a specific compartment (e.g. the ER) that would have primary consequences on local cation homeostasis. We identified that MINPP1-mediated IP_6_ regulation impacts free cations availability, as illustrated with the altered iron content of *MINPP1*^*-/-*^ HEK293 cells as well as the severe depletion of cytosolic calcium identified in *Minpp1*^*-/-*^ mouse primary neural progenitors and HEK293 cells. Interestingly, the absence of MINPP1 also severely disrupts signalling based on ER calcium that could potentially be the place of the primary defect. Calcium signaling has broad functions in neural cell physiology and brain development. Basal calcium levels influence neuronal physiology and cell survival ^45-47^, and calcium signaling plays a role in neural induction and differentiation ^48-51^. Consequently, a disruption in calcium homeostasis could be involved in PCH disease pathogenesis. A link between MINPP1 and calcium regulation has been suggested previously but it was through the synthesis of I(1,4,5)P^3 52^. Hypothetically, coupling the limitation of IP_6_-mediated chelation of calcium with the promotion of IP_3_ synthesis could be an efficient way for MINPP1 to regulate calcium signaling dynamics and homeostasis.

A mild or absent structural brain defect was also observed in other PCH mouse models. *AMPD2* null mutations cause PCH9 but the *Ampd2* single KO mouse is not associated with any obvious histological brain defect ^53^. *CLP1* is involved in tRNA processing and mutated in PCH10, however the *Clp1* mutant mouse showed only a mild decrease in the brain weight and volume, a phenotype overlooked before the identification of patients with a brain phenotype ^54^. Differences in the phenotype of human and mouse with *MINPP1* loss-of-function mutations could be related to an increased sensitivity of the human brain development to metabolic defects, although the impact of the genetic background cannot be excluded at this point.

Disrupted cation homeostasis, including metal accumulation, is central to multiple degenerative disorders^55^ such as neurodegeneration with brain iron accumulation^56,57^, Parkinson’s disease (Manganese accumulation)^58,59^, Wilson’s disease (Copper accumulation)^60,61^. Basal ganglia dysfunction is usually suspected in PCH_23_, however the severe defects identified in *MINPP1* patient MRIs suggest major neurodegeneration at the level of these subcortical nuclei, a feature not typically associated with other PCH subtypes. These structures are well known to be primarily affected by metal ions accumulation and further investigation will be needed to determine how cation chelation could contribute to the disease pathogenesis. Nevertheless, our results reveal an unappreciated basic role for highly phosphorylated IPs in cellular homeostasis which is critical during neurodevelopment.

## SUPPLEMENTARY FIGURES AND LEGENDS

**Supplementary Figure 1.**
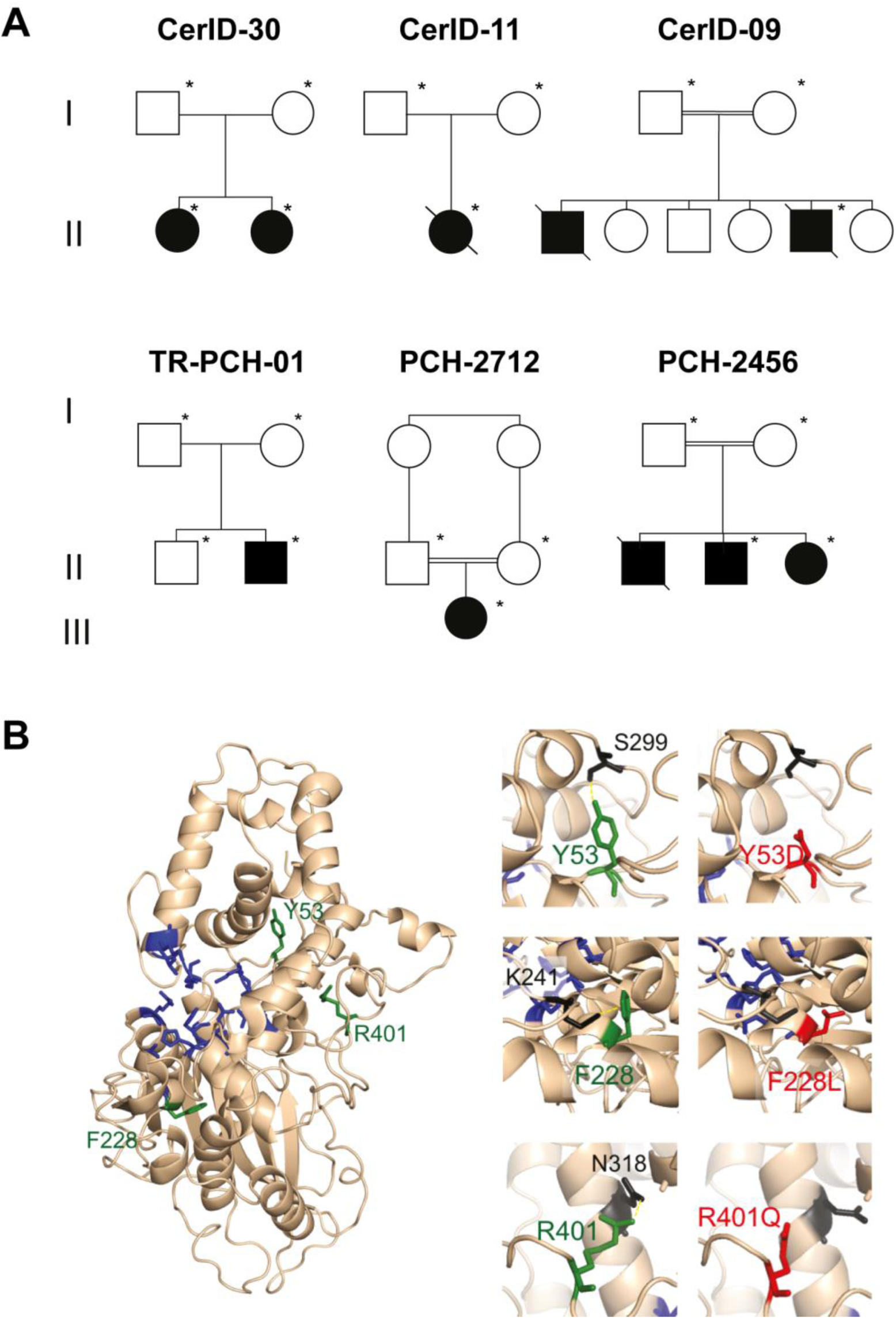
Family pedigrees and *in silico* prediction analysis of the missense *MINPP1* variants. **(A**) Pedigrees of families with mutations in the *MINPP1* gene. Individuals subjected to Whole Exome Sequencing (WES) or Sanger Sequencing are indicated with asterisk. Additional DNA samples for family CerID-09 could not be collected for segregation analysis. (**B**) Wildtype MINPP1 with amino acids involved in IP_6_ binding indicated in blue and amino acids mutated with missense variants indicated in green. The Missense3D server predicted that all the three missense variants tested cause structural damage.

**Supplementary Figure 2:**
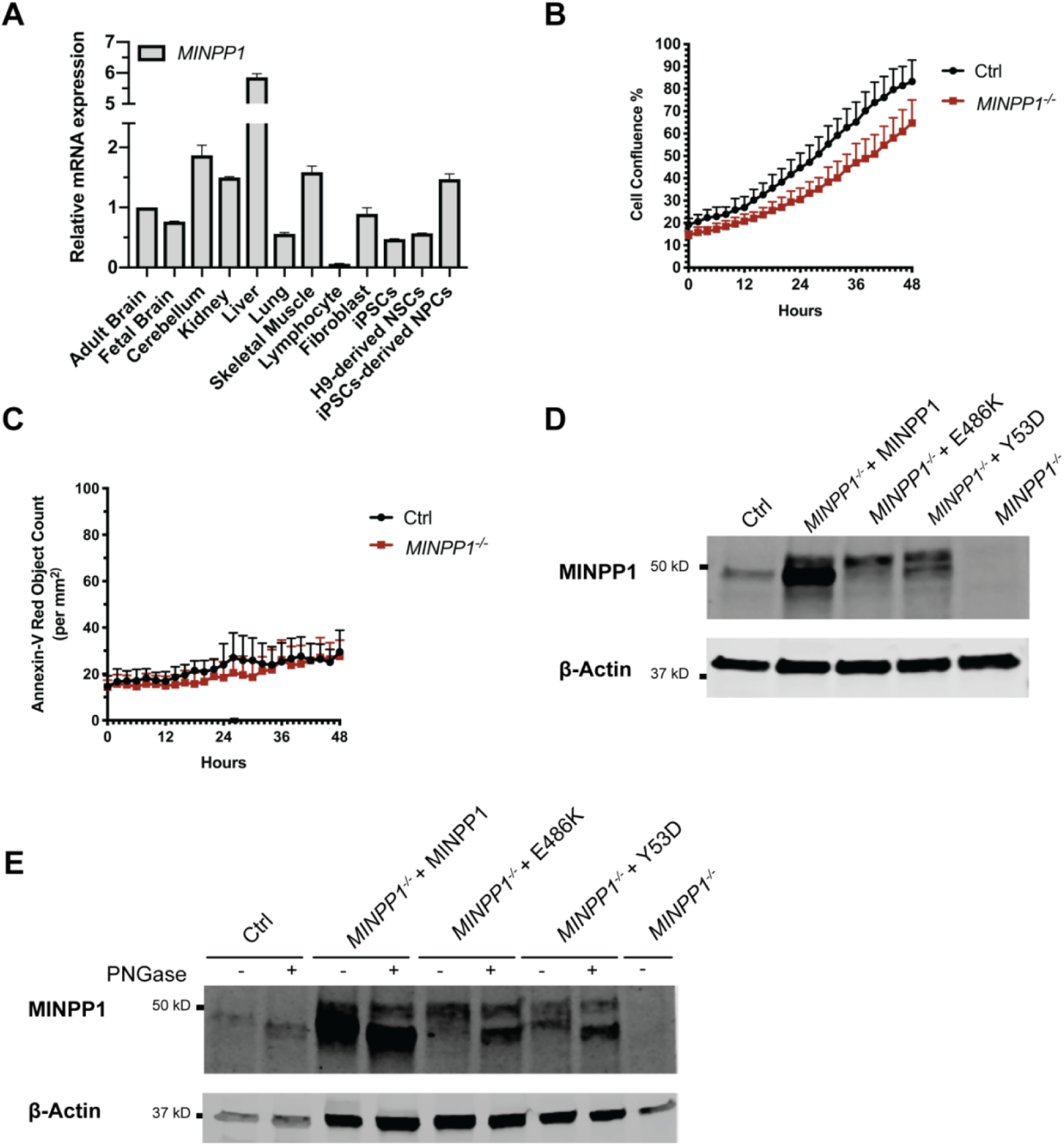
Expression profile of *MINPP1* in different human tissues and protein levels of MINPP1 variants and their corresponding glycosylation status in HEK293 cells. **(A)** Quantitative PCR analysis of human *MINPP1* expression, normalized with *ACTB*, in human cells and tissues. Abbreviations used: Neural Stem Cells (NSCs); Neuronal Progenitor Cells (NPCs) **(B)** Control and *MINPP1*^*-/-*^ HEK293 were cultured for 48 hours in IncuCyte live-cell analysis system and cell confluence (%) was analysed in IncuCyte Zoom live-cell-imaging software. Graph shows the percentage of cell confluence (± s.d.) over time. **(C)** Control and *MINPP1*^*-/-*^ HEK293 were cultured with Annexin-V containing growth media for 48 hours inside the IncuCyte live-cell analysis system, and images were captured with red fluorescence channel to detect Annexin-V positive cells. Graph shows the average ratio (±s.d.) of red cell object count per mm^2^ (n=3, two-tailed student’s t-test, not significant) **(D-E)** Western blot data showing the exogenous MINPP1 levels (D) and its glycosylated and deglycosylated forms (E) in *MINPP1*^*-/-*^ HEK293 cells transiently transfected with plasmids encoding empty vector, wild type, Y53D or E486K t MINPP1. MINPP1 is present in control HEK293 cells and absent in *MINPP1*^*-/-*^ HEK293 cells. β-Actin shown as loading control.

**Supplementary Figure 3.**
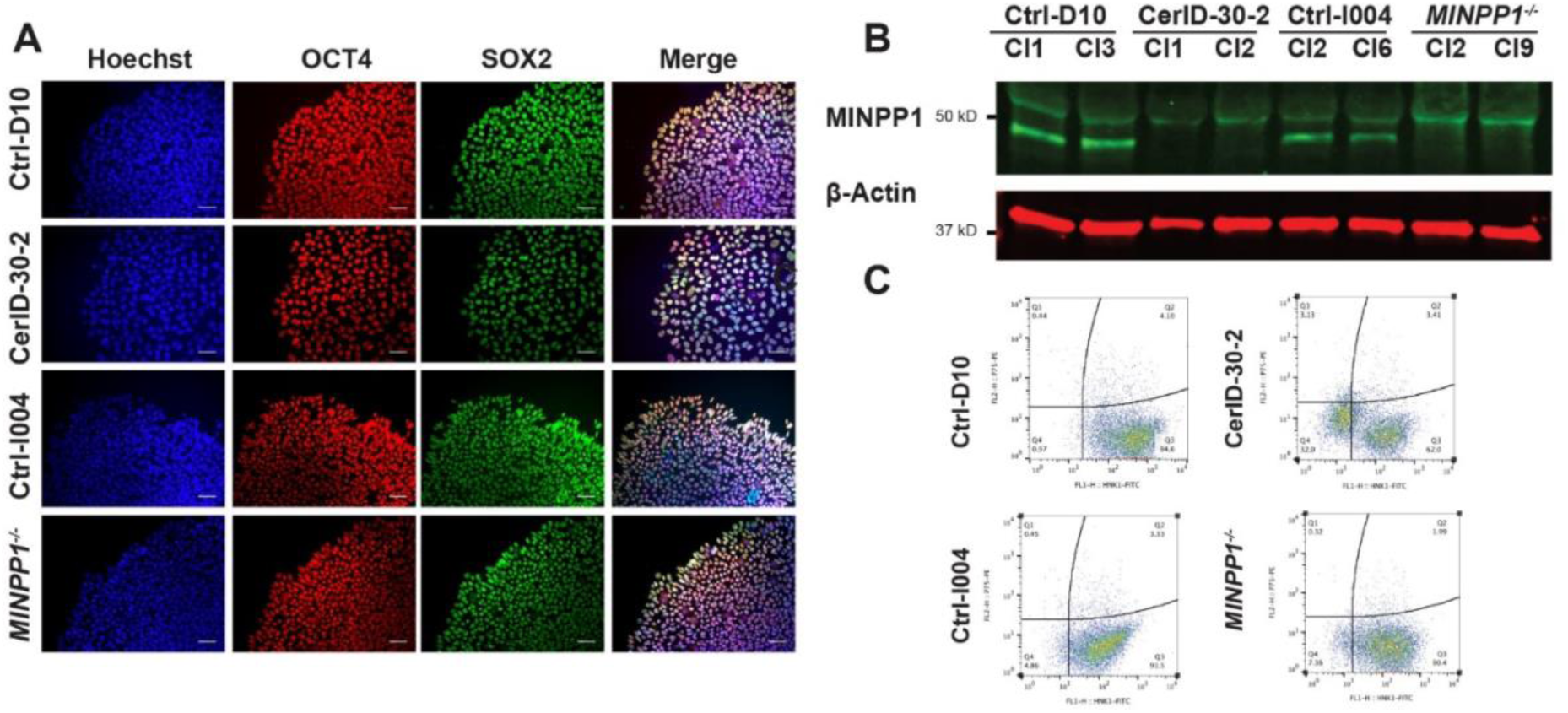
Characterization of patient-derived (CerID-30-2) and *MINPP1*^-/-^ iPSCs and their neural derivatives. **(A)** Immunofluorescence staining of embryonic stem cell markers OCT4 and SOX2 in control (Ctrl-D10 and Ctrl-I004), patient-derived (CerID-30-2), and *MINPP1*^-/-^ iPSCs. Hoechst was used as a nuclear stain. All scale bars correspond to 50μm. **(B)** Western blot data showing the absence of MINPP1 in patient-derived CerID-30-2 and MINPP1^-/-^ iPSCs. MINPP1 (48 kDa) is present in controls (D10 and I004) iPSC lines (lower band). β-actin is shown as loading control. **(C)** Flow cytometer analysis of P75 and HNK1 in control (Ctrl-D10 and Ctrl-I004), patient-derived (CerID-30-2) and *MINPP1*^-/-^ neural rosette cultures.

**Supplementary Figure 4.**
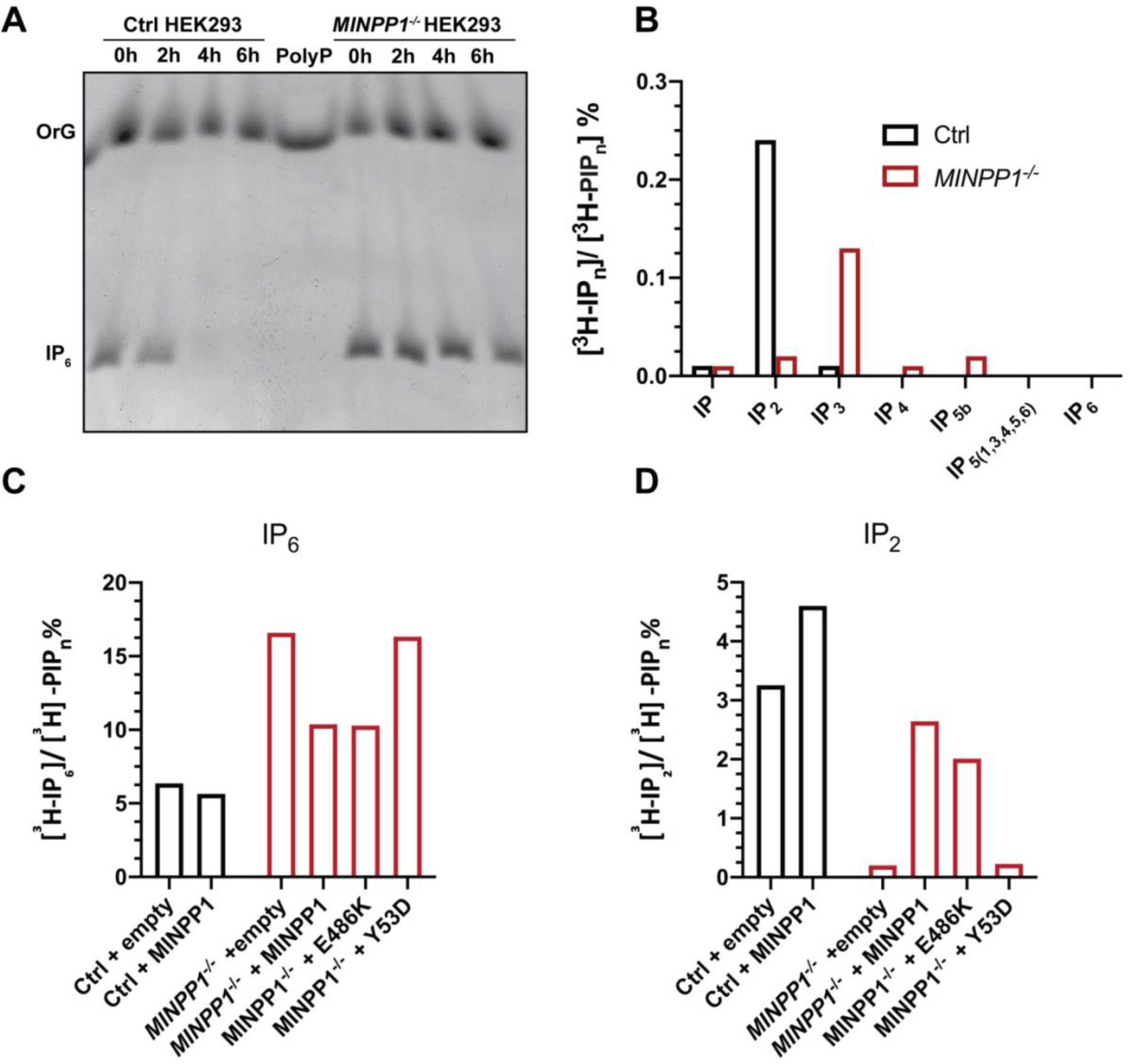
Phytase activity and SAX-HPLC analysis of extracellular and intracellular inositol phosphate levels. (**A**)MINPP1 phytase activity in conditioned medium of *MINPP1*^*-/-*^ HEK293 cells upon addition of 4 nmol of IP_6_ and incubation at 37°C for the time period indicated above. The samples were then mixed with Orange G (OrG) loading dye and resolved by PAGE followed by toluidine blue staining. Gel image representative of two independent experiments. Polyphosphate (PolyP) is shown as size ladder. (**B**) SAX-HPLC analysis of inositol phosphate levels in cell culture media of [^3^H]-inositol labeled control and *MINPP1*^*-/-*^ HEK293 cells. **(C-D)** SAX-HPLC analysis of IP_2_ and IP_6_ levels in control and *MINPP1*^*-/-*^ HEK293 cells transiently transfected with plasmids encoding empty vector, wildtype, Y53D or E486K variant MINPP1. The data is representative of two independent experiments. [^3^H]-IP_n_ levels are presented as percentage of total radioactivity in the inositol-lipid fraction ([^3^H]-PIP_n_). Abbreviations used: IPn, inositol phosphates; PIPn, phosphatidyl inositol phosphates.

**Supplementary Figure 5:**
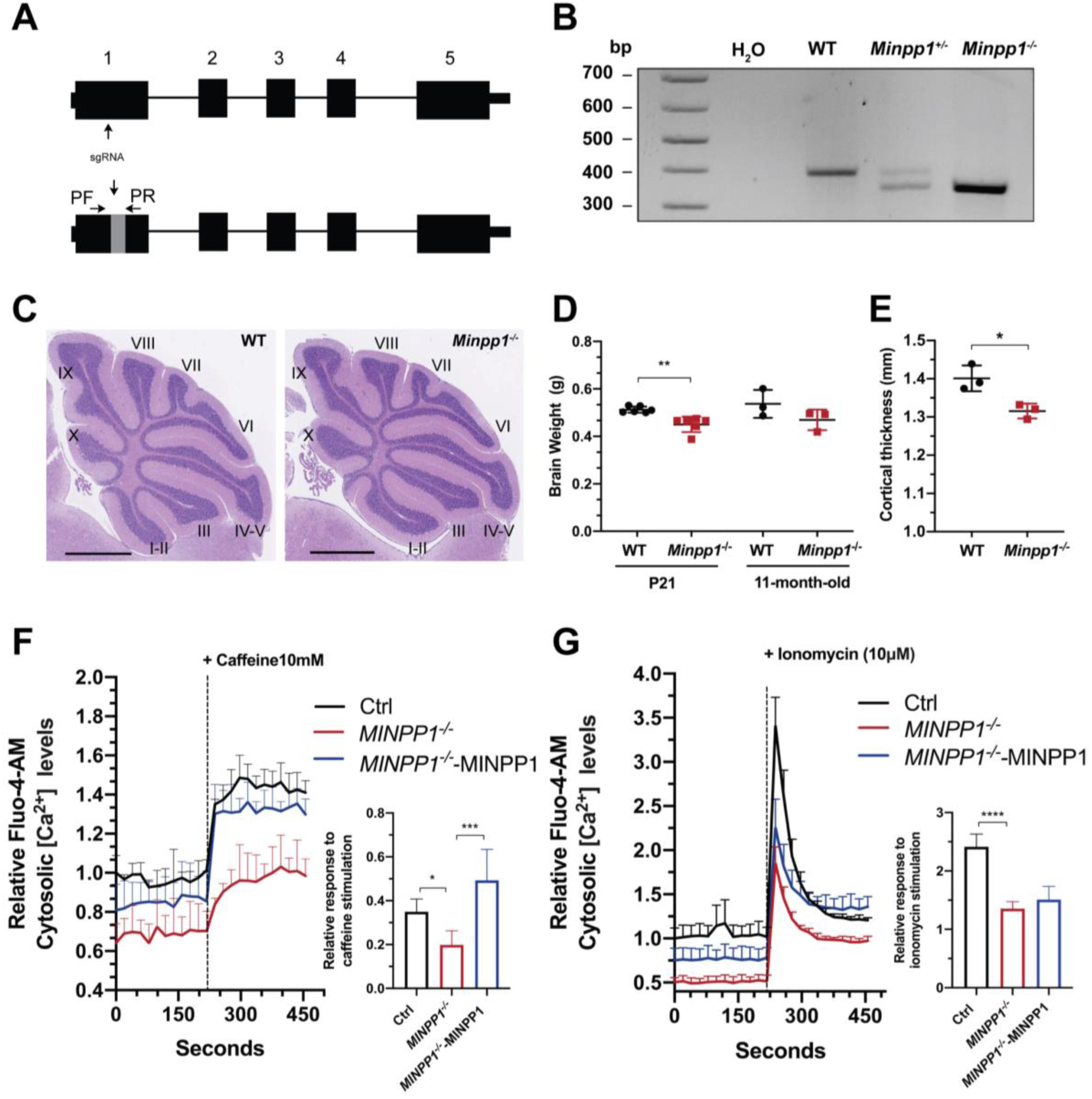
Generation and characterization of CRISPR-Cas9 Minpp1^*-/-*^ mouse and Ca^2+^ assays on HEK293 cell lines. **(A)** Schema of the sgRNA targeting site and the location of PCR primers. Abbreviations used: Primer Forward (PF) and Primer Reverse (PR). (**B**) PCR based genotyping of *Minpp1*^*-/-*^ mice indicated successful knockout alleles showing a 31 bp deletion and a smaller PCR product. (**C**) Representative images of hematoxylin-eosin (HE) staining on P21 sagittal slices of control and mutant cerebellum, scale bar 1 mm. (**D**) Quantification of total brain weight in grams (g) from P21 and 11-month old WT and *Minpp1*^-/-^ mice. (N=6 mice, Two-tailed student’s t-test, ** indicates p value p<0.01). (**E**) Quantification of cortical thickness for the WT and *Minpp1*^-/-^ cortex at P21 (N=3 mice, 30 sections per each mouse; Two-tailed Student’s t-test, * indicates p value p<0.05). (**F-G**) ) Relative Fluo-4-AM cytosolic Ca^2+^ levels in control, *MINPP1*^*-/-*^ HEK293 and MINPP1 over-expression stable *MINPP1*^-/-^ HEK293 cell lines (*MINPP1*^-/-^-MINPP1) loaded either with 10 mM caffeine (**G**) or 10 µM ionomycin in the absence of extracellular Ca^2+^. The dotted line indicates the addition of either caffeine (**F**) or ionomycin (**G**). Relative response after caffeine or ionomycin stimulation (peak) is represented graphically (inset). For all of the calcium assay experiments, the data are normalized to cell number with MTT colorimetric assay and presented as mean values relative to control baseline fluorescence intensity control ± s.d. (**F-G**) n=6, one-way ANOVA, Tukey’s post hoc test, *, ***, **** indicate p value p<0.05, p≤ 0.001 and p<0.0001 respectively).

## ACKNOWLEDGMENTS

The project is funded by the Fondation Bettencourt-Schueller-Imagine-PhD International program (EU), the French National Research Agency ANR-16-CE12-0005-01(VC), la Fondation pour la Recherche Médicale FRM-DEQ20160334938 (LC and VC) and by State funding from the Agence Nationale de la Recherche under “Investissements d’avenir” program (ANR-10-IAHU-01) and the MSDAvenir fund. A.S. and M.W. are supported by the Medical Research Council (MRC) awards (MC_UU_12018/4 and MC_UU_00012/4). We are grateful to families of patients for their participation and Dr B. Isidor for the clinical-follow up. We sincerely thank S. Jaalouk, V. Meneghini, F. Petit and A. Rotig for technical help and advice.

## AUTHOR CONTRIBUTIONS

Recruitment and evaluation of subjects: G.B., M.-T.V.-D., G.P., E.L., N.R., E.B., M.Z., M.T., F.M.S., D.M., F.G., A.M., N.B., N.B.-B., C.F. Genetic sequencing and interpretations: N.A.,V.S., C.B.-F., P.N., L.C., L.B., J.G.G., V.C. Inositol labelling and analysis: M.W. and A.S. iPSCs generation and characterization: E.U., D.M.-C., N.L. Mouse generation and characterization: K.R. and P.D. Neural differentiation and cell biology experiments: E.U. and K.R.

## ONLINE METHODS

### Patients recruitment and investigation

The patients from families CerID-09, CerID-11 and CerID-30 included in this project were referred to the Departments of Pediatric Neurology, Genetics, Metabolism or Ophthalmology of the Necker Enfants Malades Hospital. Family TR-PCH-01 was recruited by the French Reference Centre for Cerebellar Malformations and Congenital Diseases at Trousseau Hospital. Patients from families PCH-2456 and PCH-2712 were referred for neurological or genetic assessment at the National Research Center in Cairo and Hacettepe University Medical Faculty Department of Pediatric Neurology in Turkey. Written informed consents have been obtained both from the participants and the legal representatives of the children. Details regarding sequencing, filtering and prioritization protocols for each family are outlined in Supplementary Note.

### Protein Multiple Sequence Alignment

The following protein sequences are aligned through COBALT:Multiple Alignment tool: <u>NP_004888.2</u> (H. Sapiens MINPP1), NP_034929.1 (M. musculus Minpp1), NP_062136.1 (R. norvegicus Minpp1), <u>NP_957394.1</u> (D. Reiro minpp1b), NP_989975.1 (G. gallus MINPP1), XP_002935472.2 (X. tropicalis minpp1), XP_313302.4 (A. gambiae AGAP003555-PA), NP_563856.1 (A. thaliana histidine acid phosphatase family protein), NP_813655 (B. thetaiotaomicron BtMinpp)

### Maintenance and Culture of Human Induced Pluripotent Stem Cells (hiPSCs), Fibroblasts and HEK293 Cells

All human cell culture and storage protocols were performed with approval from French Research Ministry (DC 2015-2595, 09/05/2016) and all participants provided written consent. hiPSC lines (Ctrl-D10) and (CerID-30-2) were generated by Duke iPSC Share Resource Facility, using RNA-based reprogramming method. The control hiPSC line (Ctrl-I004) was generated by Imagine Institute iPSC Platform, using non-integrating Sendai virus approach (CytoTune-2.0). All hiPSC lines were shown to display embryonic pluripotency markers OCT4 and SOX2 (Supplementary Fig.3A) and no unusual chromosomal abnormalities were detected by CGH array (60K, data not shown).

hiPSCs were maintained in vitronectin-coated (10 μg/ml; 07180, STEMCELL) 3.5 cm dishes (353001, Falcon) in complete mTeSR-1 medium (STEMCELL, 85850) supplemented with 1% Penicillin-Streptomycin (PS) (15140122, Gibco) in a humidified incubator (5% CO_2_, 37 °C). The media were replaced daily and iPSCs were mechanically passaged with ROCK inhibitor Y-27632 (10 μM; 72304, STEMCELL) every 7-8 days, with a 1:3 split ratio.

Human primary fibroblasts and HEK293T (HEK293) cells were cultured in Dulbecco’s modified Eagle medium (DMEM) (11965092, Gibco) supplemented with 10% Fetal Bovine Serum (FBS) (16000044, Gibco) and 1% PS in a humidified incubator (5% CO_2_, 37 °C).

### Generation of genome edited *MINPP1*^*-/-*^ HEK293 and hiPSCs

*MINPP1*^*-/-*^ HEK293 and hiPSC lines were generated via a CRISPR-Cas9 genome editing strategy, as previously described^1^. Briefly, sgRNAs targeting the first exon of MINPP1 transcript variant 1 (NM_004897.5) were designed on CRISPOR website (http://crispor.tefor.net/) and further cloned into the pSpCas9(BB)-2A-GFP plasmid (PX458, 48138, Addgene). For the generation of *MINPP1*^*-/-*^ HEK293 clones, transfection of pSpCas9(sgRNA)-2A-GFP into HEK293 cells was performed with Lipofectamine 2000 (11668019, Invitrogen), as per manufacturer’s instructions (2.5 µg of DNA per well of a 6-well plate). Two days post-transfection, single GFP^+^ HEK293 cells were sorted into 96 well plates by Fluorescence-activated cell sorting (FACS) (BD FACSAria II SORP, BD Biosciences). Indel mutations of clones were detected by Sanger sequencing, and target editing efficiency was assessed by TIDE analysis (https://tide-calculator.nki.nl/; data not shown).

For the generation of *MINPP1*^*-/-*^ iPSCs clones, transfection of pSpCas9(sgRNA)-2A-GFP into hiPSC line (I004) was performed using Amaxa 4D-Nucleofector X-Unit (AAF-1002X, Lonza) according to manufacturer’s instructions. Briefly, hiPSCs were pre-treated with Y-27632 (10 μM) for 1 hour and further dissociated into single cell suspension with Accutase (STEMCELL, 07920). 4 x10^5^ iPSCs were nucleofected per 20 μl nucleocuvette strip, using P3 Primary Cell 4D X Kit S (V4XP-3032, Lonza), 1μg DNA per reaction, and the program CA-137. After nucleofection, iPSCs were seeded on vitronectin pre-coated 12-well plates and cultured in complete mTeSR-1 medium supplemented with Y-27632 (10 μM). Two days post-transfection, GFP^+^ iPSCs were sorted by FACS and plated at low densities of 1-2×10^4^ cells per 6cm dish (Falcon, 353037) in mTeSR-1 medium supplemented with Y-27632 (10 μM). After 12-16 days, a subset of individual colonies was processed for DNA extraction using Quick-DNA/RNA Miniprep Plus Kit (D7003, Zymo) and Sanger sequenced. Target editing efficiency was assessed by TIDE analysis (https://tide-calculator.nki.nl/; data not shown). After clonal selection and expansion, no unusual chromosomal abnormalities were detected by CGH array (60K, data not shown). Validation of *MINPP1*^*-/-*^ iPSC clones were further assessed by the absence of MINPP1 and the presence of pluripotency markers OCT4 and SOX2 with western blot and immunofluorescence staining respectively (Supplementary Fig.3A, B).

### Generation of *Minpp1*^-/-^ mice

*Minpp1*^-/-^ mice were generated with the aid of LEAT platform of Imagine Institute by using a CRISPR/Cas9 system. In this study, animals were used in compliance with the French Animal Care and Use Committee from the Paris Descartes University (APAFIS#961-201506231137361). Guide RNAs (sgRNAs) targeting the first exon of the gene were designed via the CRISPOR (http://crispor.tefor.net/). C57Bl/-J female mice (4 weeks old) were super ovulated by intraperitoneal injection of 5 IU PMSG (SYNCRO-PART^®^ PMSG 600 UI, Ceva) followed by 5 IU hCG (Chorulon 1500 UI, Intervet) at an interval of 46-48 hours and mated with C57BL/6J male mice. The next day, zygotes were collected from the oviducts and exposed to hyaluronidase (H3884, Sigma-Aldrich) to remove the cumulus cells and then placed in M2 medium (M7167, Sigma-Aldrich) into a CO_2_ incubator (5% CO_2_, 37 °C). SgRNAs were hybridized with Cas9 (Wild type) protein and injected into the pronucleus of the C57Bl/6J zygotes. Surviving zygotes were placed in KSOM medium (MR-106-D, Merck-Millipore) and cultured overnight to two-cell stage and then transferred into the oviduct of B6CBAF1 pseudo-pregnant females. The generated knockouts were validated by Sanger sequencing combined with tide TIDE analysis (https://tide-calculator.nki.nl/; data not shown). All *Minpp1*^-/-^ mice were backcrossed with C57BL/6J mice to remove potential off-targets. The *Minpp1*^-/-^ offspring were identified by Polymerase Chain Reaction (PCR) genotyping (Supplementary Fig.5). Brain histological characterization was performed with standard hematoxylin/eosin staining.

### Generation of stable HEK293 cell lines

HEK293 cells were transfected with either FLAG-HA-empty or FLAG-HA-MINPP1 plasmids by performing lipofection with the Lipofectamine 3000 reagent (Invitrogen, L3000015) as per manufacturer’s instructions (2.5 µg of DNA per well of a 6-well plate und) under serum-free media. Six hours post-transfection, the media were replaced with normal HEK293 cell culture growth media (see above). Two days-post transfection, the cell culture media were replaced with the media supplemented with the selection antibiotic geneticin (0.6mg/ml, Life, 10131027) and continued to be replaced twice a week until the cells in the 6 well plate reach high confluence. At high confluence, the cells were passaged and continued to be cultured for two to three weeks with the cell culture media supplemented with the geneticin (0.6mg/ml). The selection of clones was assessed by checking the presence of MINPP1 with Western-blot.

### E14.5 Mouse Neural Progenitor Cell Culture

Cerebral cortices (both halves) from E14.5 mice were dissected in Dulbecco Phospate Balanced Salt Solution (DPBS, 14190250, Gibco). All procedures were done at room temperature, unless otherwise stated. Cortices were then incubated for 15 minutes at 37 °C in neurocult basal medium (STEMCELL, 05700) with Trypsin (1/100, 25300062, Gibco) and DNaseI (1/1000, 79254, Qiagen). Next, neurocult basal medium with 10% FBS was added to the mixture to deactivate the Trypsin. The mixture was then titurated with fire polished Pasteur pipettes to obtain a homogenous mixture of cells. The cells were then centrifuged for 5 minutes at 300xg, washed twice and finally resuspended in neurocult medium with proliferation supplement (STEMCELL, 05701), endothelial growth factor (EGF) (20 ng/ml, PHG0314, Gibco) and 1% PS. Cells were then plated on poly-L-lysine (0.01 mg/ml, P6282, Sigma) and laminin (10 μg/ml; Gibco, 23017015) pre-coated dishes. Cells were passaged post 5 days of culture. All experiments were performed before passage 3.

### Differentiation of hiPSCs into neural lineage

hiPSCs were differentiated into neural lineage by following a published protocol^2^. In brief, hiPSCs were dissociated into small clumps with Accutase and seeded on polyornithine (0.1 mg/ml, P4957, Sigma) and laminin (4 μg/ml) pre-coated 4-well-plates (179820, Nunc) and cultured in mTSER supplemented with Y-27632 (10 μM). The day after, cell culture media were changed to N2B27 medium (1:1 mix of DMEM-F12 (31331093, Gibco) and Neural Basal Medium (1103049, Gibco)) supplemented with N2 (17502048, Gibco), B27 (17504044, Gibco), Noggin (100 ng/ml, 78060.1, STEMCELL), Y-27632 (10 μM) and 1% PS. The media were replaced every 3 days.

### Inositol Labelling and SAX-HPLC Analysis

HEK293 cells and fibroblasts were plated at 10,000 cells/well on 6-well plates and radiolabeled for 5 days with 5 μCi/ml *myo*-[^3^H]-inositol (1 mCi, NET114A001, Perkin Elmer) in inositol-free DMEM medium (DML13-500, Caisson) supplemented with dialyzed FBS (Gibco,26400-044) and 1%PS. One quarter of the media was replaced with fresh medium containing 5 μCi/ml [^3^H]-inositol on day 2. For iPSCs and their day 7 neurally-differentiating counterparts, the inositol metabolic radiolabeling was performed for 3 days and inositol-free DMEM media were supplemented with either bFGF (10 ng/ml, PHG0264, Invitrogen) or Noggin (100 ng / ml, 78060.1, STEMCELL) respectively. The latter time point was chosen to analyze the inositol phosphate levels at day 10 differentiating stage, the time point corresponding to neuronal differentiation stage shortly after neural induction and before the observation of differences in the TUJ1/PAX6 cell populations between control and *MINPP1* mutant iPSCs. For in vitro overexpression studies, the inositol metabolic radiolabeling was performed for 3 days, followed by lipofectamine-based transfection into HEK293 cells in the presence of 5 μCi/ml *myo*-[^3^H]-inositol. One day post-transfection, the inositol phosphates were extracted. Extraction of inositol phosphates was performed according to published protocol^3^. Briefly, cells were incubated on ice with 1 M perchloric acid and 5 mM EDTA for 10 minutes followed by the collection of extracts and an overnight neutralisation step with 1 M potassium carbonate. Inositol phosphates were then separated by strong anion-exchange high-performance liquid chromatography (SAX-HPLC). The peaks ([^3^H]-IP^n^) were identified based on comparison to standards according to a previously published study^3^. The radioactivity within each fraction was measured with a beta counter after addition of 4 ml of scintillation fluid (Ultima-Flo AP LCS-mixture; Packard).The data are presented as percentage of total radioactivity in the inositol-lipid fraction, obtained by incubating the post-extraction cells with 0.1% NaOH and 0.1% SDS overnight at room temperature.

### Cloning of plasmids

The MINPP1 cDNA (transcript NM_004897.5) sequence was isolated by PCR from human brain cDNA. The PCR product was then ligated into the FLAG-HA-pcDNA3.1 vector (52535, Addgene) between XhoI and BamHI sites. FLAG-HA-pcDNA3.1 was a gift from Adam Antebi (Addgene plasmid # 52535; http://n2t.net/addgene:52535; RRID: Addgene_52535). The mutations c.157T>G and c.1456G>A were incorporated in to the wild type plasmid with QuickChange Lightning Site-Directed Mutagenesis Kit (210515, Agilent). One Shot TOP10 chemically competent *E. coli* (C404006, Invitrogen) were then transformed with the resulting plasmid and plated on ampicillin resistant agar plates. Plates were incubated at 37 °C overnight. Colony PCR was performed on plasmids by using T7 promoter/BGH-rev primers and clones were sequenced by Sanger sequencing to check sequence reliability.

### Transfection experiments for proliferation studies using HEK293 cells

Transfection into HEK293 cells was performed using Amaxa 4D-Nucleofector X-Unit (AAF-1002X, Lonza) according to manufacturer’s instructions. In brief, 2 x10^5^ HEK293 cells were nucleofected per 20μl nucleocuvette strip, using P3 Primary Cell 4D X Kit S (V4XP-3032, Lonza), 1μg DNA per reaction, and the program CA-137.

### MTT Proliferation Assay

Cells were cultured with Methylthiazolyldiphenyl-tetrazolium bromide (MTT) (0.5 mg/ml, M5655, Sigma) for 1.5 hours. Then, MTT and the cell culture medium were removed and 100% dimethyl sulfoxide (100 μl/well) was added to dissolve the formazan crystals. Next, the cell culture plate was left to shake for 15-30 minutes in the dark at room temperature, followed by transfer of the samples into flat-bottom 96-well microtiter plate (655101, Greiner). The optical density was read on a microplate reader at 560 nm (1681135, Biorad).

### Quantification of free intracellular Ca^2+^ levels

Cells were plated at 25,000 cells/well on flat-bottom 96-well plates (Greiner, 655088). After 24 to 48 hours in culture, the cells were loaded with Fluo-4-AM (5 mM, F14201, Invitrogen) diluted in assay buffer (NaCl 140 mM, Glucose 11.5 mM, KCl 5.9 mM, MgCl_2_ 1.4 mM, NaH_2_PO_4_ 1.2 mM, NaHCO_3_ 5 mM,, HEPES 10 mM, pH 7.4) with or without CaCl_2_ (1.8 mM) to a final concentration of 5 μM. After 30 minutes of incubation in a humidified incubator (5% CO_2_, 37 °C), the loading medium was removed and the cells were washed twice with the assay buffer. Then, the cells were excited at 485nm and emission intensities was recorded at 535nm via a fluorometers (200ProTecan and Tristar LB941) in the presence of assay buffer. The measured fluorescence intensities were then normalised to cell number via MTT proliferation assay. For Ca^2+^ stimulation experiments, Ionomycin (10 μM, I0634-1MG, Sigma) or Caffeine (10 mM, C0750, Sigma) were quickly added to the wells in the presence of assay buffer and fluorescence intensities were recorded every 30 seconds for a minimum of 10 minutes. All the Ca^2+^ stimulation experiments were performed with early batch HEK293 cells^4^.

### Quantification of intracellular total and free iron levels

The total iron content was measured with a colorimetric ferrozine-based assay according to previous studies^5,6^. Briefly, cells were cultured on 6-well plate (92006, TPP) and incubated with or without ferric ammonium citrate (FAC) (100 μM, RES20400-A702X, Sigma) in serum-free DMEM for 48 hours. Then, the cells were lysed with NaOH (50 mM, 200μl/well) and 50 μl of the lysate was kept apart for protein quantification. The remaining cell lysate were mixed with equal volumes of 10 mM HCl and iron-releasing reagent (HCl 1.4 M, KMnO_4_ 4.5%) and incubated for 2 hours at 60 °C. Once the samples reached room temperature, 30 μl of iron-detection reagent (6.5 mM ferrozine, 6.5 mM neocuproine, 2.5 M ammonium acetate, and 1 M ascorbic acid) was added to each sample. After 30 minutes of incubation at room temperature, the samples were transferred to flat-bottom 96-well microtiter plate (Greiner, 655101). The optical density was read on a microplate reader at 560 nm (Biorad, 1681135). FAC was used as reference standard.

The free iron levels (Fe^2+^ and Fe^3+^) were measured by using the Iron Assay Kit (Abcam, ab83366) according to manufacturer’s instructions. In both iron-detection assays, the data are normalized against the protein levels, quantified via Pierce BCA Protein assay kit (23225, Thermo Scientific).

### Immunofluorescence

Cells were fixed in cold 4% paraformaldehyde for 10 minutes followed by three washes of PBS and permeabilization in 0.25% PBS-Triton-X-100 for 10 minutes. Cells were blocked for 1 hour in 1% BSA and 22.52 mg/ml glycine diluted in 0.1% PBS-Tween-20 (PBST). Cells were then incubated overnight at 4 °C with the following primary antibodies diluted in 1% BSA/0.1% PBST: Anti-PAX6(1:400, 901301, BioLegend), Anti-TUBULIN β-3 (1:1000, 801201, BioLegend), Anti-OCT4 (1:100, sc-5279, Santa-Cruz), Anti-SOX2 (1:1000,AB5603, Millipore). After 3 washes with 0.1% PBST, cells were incubated for 1 hour with Alexa-Fluor-coupled secondary antibodies (A21424, A-11034, Invitrogen) diluted 1:500 in 0.1% PBST. Following three washes with 0.1% PBST, cells were then incubated with Hoechst 33342 (H3570, Invitrogen) diluted 1:1000 in 0.1% PBST. After 10 minutes of incubation, cells were washed twice with 0.1% PBST and either kept in PBS or mounted with ProLong™ Diamond Antifade Mountant (P36965, Invitrogen). Images were acquired with CELENA-S Digital Imaging System (CS20001, Logos) and Zeiss Axioplan-2. For quantification analysis, a minumum of 10 fields per replicate were analyzed in ImageJ. The percentage of PAX6^+^ or TUJ1^+^ cells were counted relative to the total number of Hoecsht^+^ cells per field which was comparable (i.e., Hoecsht^+^) between replicates and conditions.

### Protein extraction and western blot analysis

Protein extraction, quantification, separation in gel electrophoresis and transfer were performed as described previously^1^. Nitrocellulose membranes were blocked either in Odyssey-TM Blocking Buffer (927-50003, LICOR) or in 5% dry milk diluted in 0.2% PBST for 1 hour and further incubated overnight at 4 °C with the following primary antibodies diluted in either OdysseyTM Blocking Buffer or 2.5% dry milk diluted in 0.2% PBST: Anti-MINPP1 (1:2000, sc-10399, Santa-Cruz), Anti-β Actin (1:5000, AM4302, Invitrogen), Anti-Calmodulin (1:1000, 465, Swant). After three washes with 0.2% PBST, the membranes were then incubated for 1 hour at room temperature with HRP (1:10000, sc-2314, sc-2020, Santa-Cruz) or IRDye-coupled (1:10000, 925-68070, 925-32211, LICOR) secondary antibodies. After three washes with 0.2% PBST, the membranes were developed either with HRP (ChemiDoc XRS, Biorad) or Odyssey CLx imaging system (LICOR).

### Statistics

Data were analysed where appropriate with one-way analysis of variance (ANOVA), Two-tailed student’s t-test and two-way ANOVA with GraphPad Prism 8. N and n values are detailed in figure legends. “N” refers to independent experiments or animals); “n” refers to technical replicates.

All the data are presented as mean ± the standard deviation (s.d.). Statistically significant differences are indicated on the figures by *p<0.05, **p<0.01, ***p<0.001, or ****p<0.0001.

## Methods for Supplementary Figures

### Construction of human MINPP1 structural model

The human MINPP1 structure model was generated with the protein structure homology-modeling server Phyre V2.0 (PMID: 25950237) based on the D. castellii phytase structure (PDB: 2GFI) with 89% sequence coverage. The impact of the missense mutations involving amino-acids included in the MINPP1 structure (i.e. Tyr53, Arg401, Phe228) was evaluated using Missense3D (http://www.sbg.bio.ic.ac.uk/~missense3d/)^7^. All the structural figures were prepared with Pymol (http://www.pymol.org).

### RNA isolation, RT-qPCR and relative fold change expression analysis

The total RNAs of the different tissues used in this study were ordered from Clontech Laboratories (USA); Human brain (636530); Human fetal brain (636526); Human cerebellum (636535); Human liver (636531); Human lung (636524); Human kidney (636529) and Human skeletal muscle (636534). Total RNA extraction was performed from cell culture pellet using TriZol (ThermoFischer, 15596026) according to the supplier’s instructions. The purification of the RNA was done with the RNeasy Mini Kit (Qiagen, 74104). Reverse transcription of the total RNAs was performed using SuperScript II reverse transcriptase (ThermoFisher, 18064022) according to the manufacturer’s recommendations. Quantitative PCR was performed with the SYBR Green PCR Master Mix reagent (ThermoFisher, 4364346) on an Applied Biosystems One Step Plus real-time PCR system (Applied Biosystems, 4376600.*ACTB* was chosen as a reference gene to normalize the results between different tissues and cell line. Primers used for qPCR were: CCCTTGCCATCCTAAAAGCC (Forward) and TGCTATCACCTCCCCTGTGT (Reverse) for *ACTB*; CCCTTGCCATCCTAAAAGCC (Forward) and TGCTATCACCTCCCCTGTGT (Reverse) for *MINPP1*. The relative expression of *MINPP1* was determined by the ΔΔCt method using the human brain as the reference tissue.

### Glycosylation Studies

The glycosylation status of different MINPP1 mutant forms is checked by following the previously described PnGase F protocol (P0704L, New England BioLabs) ^2^.

### Cell Proliferation and Apoptosis Assay with Incucyte-Live-Imaging System

Cells were seeded on 96-well plate (4379, Essen Bioscience) and cultured with Annexin-V (1:200, 4641, Essen Bioscience) containing media inside the IncuCyte® Live Cell Analysis System (Essen Bioscience). Three images per well were taken with phase contrast and red fluorescence channels every 2 hours for a total of 48 hours, using 20x objective. Images were then analysed with Incucyte-Zoom software, by defining a mask with the basic analyzer. The same mask was applied to all time points.

### Differentiation of hiPSCs towards dorsal telencephalic progenitors

hiPSCs were differentiated towards dorsal telencephalic lineage by following the protocol as previously described ^1,8^. Briefly, hiPSCs were manually dissociated into big clumps and floated in non-coated dishes in neural induction media N2B27 (1:1 mix of DMEM-F12 (31331093, Gibco) and Neural Basal Medium (1103049, Gibco)) supplemented with N2 (07156, STEMCELL), NeuroCult SM1 without vitamin-A (05731, STEMCELL), bFGF (10 ng/ml, PHG0264, Invitrogen) and dual-SMAD inhibition small molecules SB431542 (20 µM, 72232, STEMCELL) and LDN193189 (HCl) (500 nM, 72146, STEMCELL). After 6 hours, embryoid body (EB)-like structures were seeded on polyornithine (0.1 mg/ml, P4957, Sigma) and laminin (4 μg/ml) pre-coated 3.5 cm dishes. The media were changed every 3 days. At day 12-15, the neural rosettes were manually or enzymatically passaged on to polyornithine-laminin-pre-coated dishes and cultured in N2B27 medium supplemented with bFGF (10 ng/ml), EGF (10 ng/ml, PHG0314, Invitrogen) and BDNF (20 ng/ml, 78005, STEMCELL). At confluence, cells were passaged with a density of 5×10^4^/cm^2^ and the media was supplemented with Y-27632 (10μM) on the day of passage. Neuroectodermal origin of the emerging neural progenitor-like cells was assessed by HNK1/P75 Flow cytometry (Supplementary Fig.3C).

### Flow Cytometry

Cells were fixed with 1% paraformaldehyde and cell suspensions in PBS were stained for 30 minutes with anti-HNK1-FITC (130-092-174, MiltenyiBiotec), anti-P75-APC (130-110-079, MiltenyiBiotec) REA control-FITC (130-113-449, MiltenyiBiotec), or REA Control -PE (130-113-450, MiltenyiBiotec) at 1/10 dilution. Prior to analysis, the samples were centrifuged and resuspended in PBS. The samples were then processed with FACSCalibur (BD Biosciences) and analyzed with FlowJo Software (Treestar, Ashland, OR).

### *In vitro* phytase Assay

The *in vitro* phytase assay was performed as previously described^9^. Briefly, 4 nmol of IP_6_ was added to 1 ml of conditioned media from Ctrl and *MINPP1*^-/-^ HEK293 cells and further incubated for 0, 2, 4 and 6 hours at 37 °C. The samples were then mixed with Orange G loading dye and resolved by 35% PAGE followed by toluidine blue staining.

### Analysis of extracellular inositol phosphates

For analysis of extracellular inositol phosphates, cells were labelled for 5 days with [3H]-inositol as indicated above. Media were collected and centrifuged at 300xg for 5 minutes to remove cell contaminants. Perchloric acid was added to the supernatant to final 1 M. The acidified medium was incubated on ice for 10 minutes before centrifugation at 18,000 xg to remove precipitates. Inositol phosphates in the supernatant were then purified using titanium dioxide beads^10^. Briefly, 4 mg beads were added to the media extracts and the samples rotated at 4°C for 15 minutes. Beads were washed twice in 1 M perchloric acid, before elution with ammonium hydroxide. Radiolabeled inositol phosphates in the purified media extracts were separated by SAX-HPLC as before, and radioactivity in each fraction measured with a beta counter.

## Supplementary Note

### Detailed method for Sequencing, Variant Filtering and Prioritization

#### Families CerID-09, 11 and 30

Genomic DNA was extracted from peripheral blood cells. As previously published (Chemin et al. Brain. 2018; Megahead et al. Orphanet J Rare Dis. 2016), Exome capture was performed with the the SureSelect Human All Exon kit (Agilent Technologies). Agilent Sure Select Human All Exon (54Mb Clinical Research Exome) libraries were prepared from 3µg of genomic DNA sheared with an Ultrasonicator (Covaris) as recommended by the manufacturer. Barcoded exome libraries were pooled and sequenced with a HiSeq2500 system (Illumina), generating paired-end reads. The mean depth of coverage of the exome libraries was greater than ∼100-120X with >98 to 99% of the targeted exonic bases covered at least by 15 independent reads and >93 to 98% by at least 30 independent sequencing reads (98-99% at 15X and 93 to 98% at 30X). After demultiplexing, sequences were aligned to the reference human genome hg19 using the Burrows-Wheeler Aligner (BWA). Downstream processing was carried out with the Genome Analysis Toolkit (GATK), SAMtools and Picard, following documented best practices (http://www.broadinstitute.org/gatk/guide/topic?name=best-practices). Variant calls were made with the GATK HaplotypeCaller (V3.7). The annotation process was based on the latest release of the Ensembl database, Gnomad 2.1. Variants were annotated and analyzed using the Polyweb software interface designed by the Bioinformatics platform of University Paris Descartes. Variants were filtered to less than 1% minor allele frequency in population databases and with MAF <0.001 in our sequenced cohort that include more than 15000 exomes. Manual curation of the known PCH genes (for the gene list see Chemin et al. Brain 2018) was first performed to exclude a known genetic cause. Variants were then prioritized by the suspected mode of inheritance, OMIM status and *in silico* prediction tools (Mutation Taster (Schwarz et al. Nature Methods. 2014); SIFT and Polyphen (McLaren et al. Genome Biology. 2016); CADD score (Rentzsch et al. Nucleic Acids Res. 2018)). Candidate genes were correlated to the patient’s phenotype, gene function and expression. Sanger sequencing of the patient and her parents was performed to confirm the variants and co-segregation using a 3500xl genetic analyzer (4405633; Applied Biosystems). For the family CerID-09, homozygous variants were identified in 31 genes. From these analyses, 3 genes with the predicted most likely pathogenic variants (*MINPP1*; *ZCCHC24* and *NKX1-2*) were further considered as candidates. For the family CerID-11, 4 variants were identified in 2 potential candidate genes with 2 heterozygous variants (*MINNP1* ; *KIAA1522*). For the family CerID-30, 21 variants were identified in 11 potential candidate genes including 6 genes with 2 or more heterozygous variants, and 5 genes with one homozygous variant. Two genes with most likely pathogenic variants were further explored: *MINPP1* and *MTFMT*.

#### Family TR-PCH-01

Patient TR-PCH-01-1 was part of a replication cohort of 100 patients with congenital or early-onset cerebellar defects that includes 21 PCH cases. This cohort was investigated using a targeted gene panel. This panel includes 184 known genes involved in congenital cerebellar anomalies (list available upon request), and 13 candidate genes including *MINPP1*. Samples were prepared using Blood DNA and the SeqCap EZ Choice preparation kit (Roche) and sequenced on an Illumina MiSeq sequencer using 2 × 150 bp sequencing kits. The Basespace cloud computing platform (with BWA 2.1 and GATK Unified Genotyper 1.6) and the Variant Studio v.3.0 software provided by Illumina were used. Other known PCH genes were excluded for TR-PCH1-1. Pathogenicity of variants was ascertained according to the ACMG (American College of Medical Genetics) criteria. The segregation was confirmed by Sanger sequencing in the index case, both parents and unaffected sibling.

#### Families PCH-2456 and PCH-2712

Blood DNA was extracted using Qiagen reagents (Qiagen Inc., USA) and whole exome sequencing was performed on all affected family members using the Illumina HiSeq2500 which yields 100 bp paired-end reads covering 80% of the exome at 20X. GATK best practices pipeline was used for SNP and INDEL variant identification. Variants were prioritized by allele frequency (<0.1% MAF in our exome database of over 5000 individuals), conservation and predicted effect on protein function. The variants were predicted to be disease causing and were absent from 1000 Genomes, ExAC and gnomAD. Sanger sequencing confirmed segregation according to a strict recessive mode of inheritance in all genetically informative and available family members.

